# A novel polyubiquitin chain linkage formed by viral Ubiquitin prevents cleavage by deubiquitinating enzymes

**DOI:** 10.1101/756791

**Authors:** Hitendra Negi, Pothula Puroshotham Reddy, Chhaya Patole, Ranabir Das

**Affiliations:** National Center for Biological Sciences, TIFR, Bangalore, India; SASTRA University, Thirumalaisamudram, Thanjavur, India; Institute for Stem Cell Science and Regenerative Medicine, Bangalore, India

**Keywords:** Nuclear Magnetic Resonance, Virus, Ubiquitin, Post-translational Modifications, Host-pathogen interactions

## Abstract

The *Baculoviridae* family of viruses encode a viral Ubiquitin gene. Although the viral Ubiquitin is homologous to eukaryotic Ubiquitin (Ub), preservation of this gene in the viral genome indicates a unique function that is absent in the host eukaryotic Ub. We report the structural, biophysical, and biochemical properties of the viral Ubiquitin from *Autographa Californica Multiple Nucleo-Polyhedrosis Virus* (AcMNPV). The structure of viral Ubiquitin (vUb) differs from Ub in the packing of the central helix α1 to the beta-sheet of the β-grasp fold. Consequently, the stability of the fold is lower in vUb compared to Ub. However, the surface properties, ubiquitination activity, and the interaction with Ubiquitin binding domains are similar between vUb and Ub. Interestingly, vUb forms atypical polyubiquitin chain linked by lysine at the 54^th^ position (K54). The K54-linked polyubiquitin chains are neither effectively cleaved by deubiquitinating enzymes, nor are they targeted by proteasomal degradation. We propose that modification of proteins with the viral Ubiquitin is a mechanism to counter the host antiviral responses.

## INTRODUCTION

Ubiquitin (Ub), a 76 residue protein, regulates multiple cellular pathways and is a major way of regulating protein levels by the Ubiquitin-proteasome pathway (1–3). Ubiquitin is covalently attached to the target substrate *via* an isopeptide bond to generate a post-translation modification (PTM). The process of conjugating Ubiquitin to different substrates is a three-step process involving enzymes *viz.* Ubiquitin-activating enzyme E1, Ubiquitin-conjugating enzymes E2s, and finally, Ubiquitin ligase enzymes E3s. At the end of conjugation, the C-terminal glycine residue of Ubiquitin forms an isopeptide bond with the ε-amino group of the substrate’s lysine residue. Furthermore, Ubiquitin can build polyubiquitin chains via seven lysine residues, and depending on chain linkage, the function of the polyUb chain is decided (4). For e.g., a substrate with K63 polyUb chains is involved in DNA repair pathways (5) while K11 and K48 polyUb chains eliminate the substrate *via* proteasomal degradation (6).

The Ubiquitin pathway plays a major role in host-virus interactions (7). Often viruses co-opt the host ubiquitination machinery for effective infection (8). In some cases, viruses also encode genes that express enzymes involved in ubiquitination reaction (9). These enzymes help the virus to degrade host immune response, avoid detection during latency, enhance viral genome transcription/replication, and facilitate egress (10, 11). Baculoviridae family of viruses infect a variety of invertebrates such as Arthropods, Lepidoptera, Hymenoptera, Diptera, etc. thus commonly referred to as insect viruses. Alphabaculovirus is the largest genera of this virus family, which contains almost all the 33 Nuclear Polyhedrosis viruses (NPVs). The characteristic feature of NPVs is to survive harsh and non-conducive environments by the formation of an extremely stable polyhedral capsid, which later turns to protein crystal (12). *Autographa californica Multiple Nucleo-Polyhedrosis Virus* (AcMNPV) is the most studied virus of this genus containing a 134 Kbp double-stranded genome with 154 ORFs packaged into rod-shaped nucleocapsids (13). Interestingly, a large number of NPV viruses, including AcMNPV, express an Ubiquitin-like molecule known as viral Ubiquitin (vUb).

Eukaryotic Ubiquitin has 76% sequence identity with vUb (14). Although the presence of Ubiquitin within the baculovirus genome likely arose by horizontal gene transfer, its retention suggests that vUb provides some specific advantage, which is not provided by the host eukaryotic Ub. vUb expresses with viral coat proteins in the late phase of the virus life cycle (15). Deletion of the vUb gene reduces the generation of virions, suggesting its role in forming or assembling viral particles (16). Recently, it was discovered that vUb plays an important role in specifying whether a nucleocapsid will form a budded virus or occlusion-derived virus (17). Other postulations of vUb function include proteasomal degradation of coat proteins for uncoating the viral capsid during early stages of infection. However, the structural details of vUb are unavailable, and it is unknown if vUb interacts with the proteasome. Apart from the seven conserved lysines present in eukaryotic Ubiquitin, vUb has an extra lysine at the 54^th^ position (K54). Whether vUb can form atypical K54-linked polyubiquitin chain, and the functional role of such a chain is also unknown.

Here we report the solution structure of vUb. While the overall fold of Ub is conserved in vUb, there are differences in core packing between the two proteins. A comparison of the stability between Ub and vUb confirms that differences in core packing reduce the stability of vUb. The interaction of vUb with the Ubiquitin binding domains in proteasome receptors was studied, which suggested that the vUb binds equally well to these receptors as Ub. The efficiency of polyubiquitination was comparable between Ub and vUb. Finally, we confirmed that vUb assembles K54-specific polyubiquitin chains. These chains are not effectively cleaved by the host DUBs. The K54-specific chains are also not targeted for proteasomal degradation. Our study proposes that the virus encodes vUb to conjugate substrates with the unique linkage of polyubiquitin chain. Conjugation of viral protein substrates by such polyubiquitin chain may provide a mechanism to avoid antiviral responses that regulate the conjugated substrates.

## Results and Discussion

### Sequence conservation of viral ubiquitin

A gene tree was reconstructed using Maximum Likelihood method from all Ubiquitin coding amino-acid sequences taken from 39 baculoviruses, out of which 31 are *Nucleo Polyhedrosis Virus* (NPV) including AcMNPV and 8 are Granulovirus (GV) (Table S1). The gene tree suggests all the GVs are monophyletic and NPV and MNPVs are polyphyletic groups. Interestingly, all the K54 containing viruses share a common ancestry, except EpNPV, which may have undergone parallel evolution (Fig 1). The Ubiquitin surface is generally polar, with the exception of a large hydrophobic region centered near the C-terminal end of β strand 5 that includes Leu8, Ile44, and Val70. The hydrophobic patch plays a role in many interactions with Ubiquitin binding domains and enzymes in the Ubiquitination reaction. The multiple sequence alignment suggested that the hydrophobic surface patch is conserved in viral Ubiquitin sequences. On the contrary, the buried aliphatic hydrophobic residues like Leu15, Ile23, and Val26 are not necessarily conserved. By the stability-function tradeoff, vUb may have diverged to include other functionalities at the cost of protein stability.

**Figure 1.**
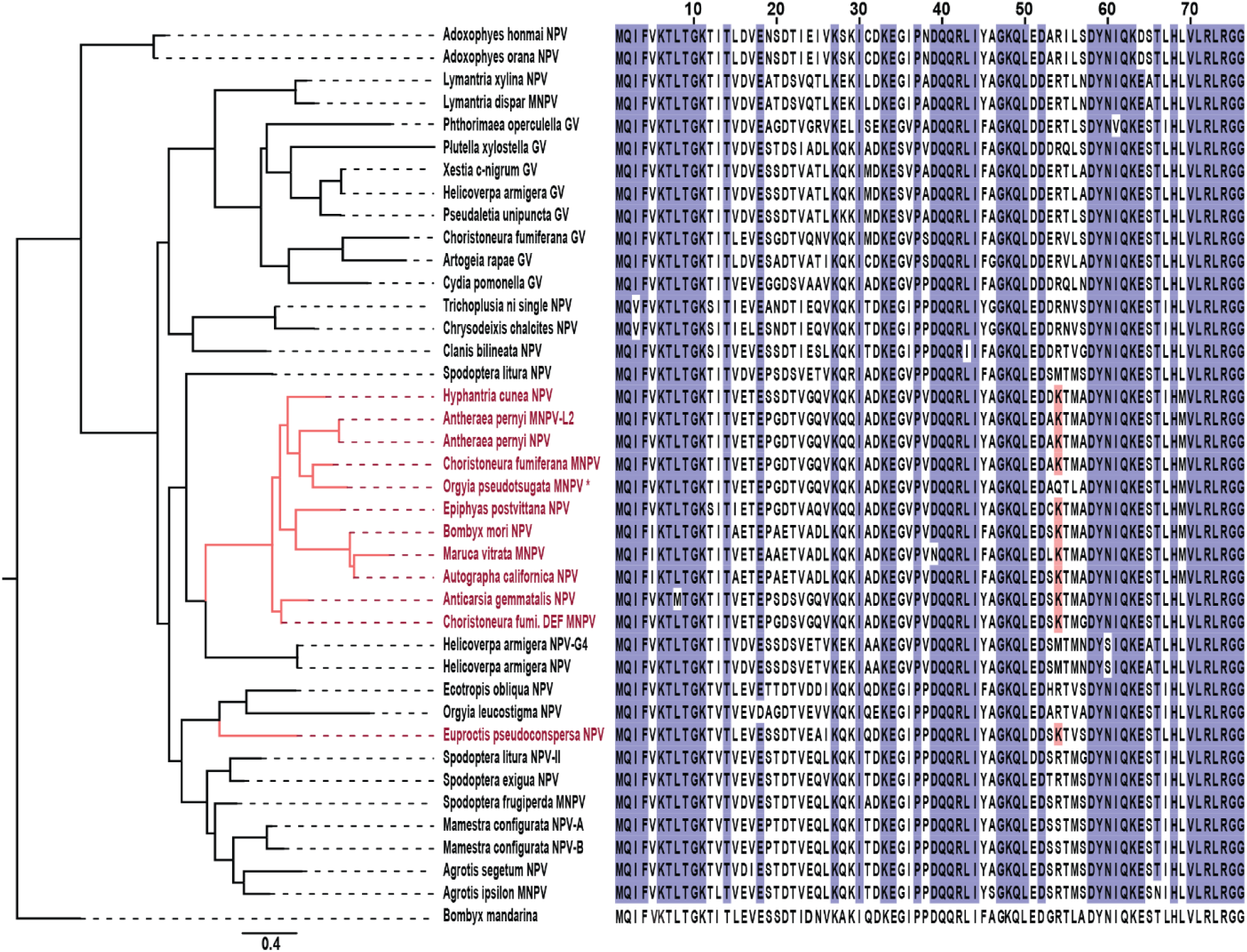
Sequence analysis of vUb. Maximum Likelihood tree of vUb among 39 insect viruses (31 Nuclear Polyhedrosis Virus and 8 Granuloviruses) based on the amino acid sequences. All the insect viruses with extra lysine at 54^th^ position are highlighted in red. Ub sequence from *Bombyx Mandarina* (host) is used as an outgroup.

### Solution structure of viral ubiquitin

The sequence identity of vUb from AcMNPV with human Ubiquitin (Ub) is 76% (Fig 2A). The canonical hydrophobic patch (L8, I44, and V70) that is crucial for activation of Ub by E1 and E2 enzymes is conserved. The seven lysines that are used to form polyubiquitin chains are also conserved. The major differences in the sequence are observed in helix α1, the loop between β2 and α1, and the loop between β4 and β5. The structure of vUb was determined to see if these differences in the sequence have any structural implications for vUb. Uniformly double-labeled (^13^C, ^15^N) vUb sample was prepared to determine its solution structure. The sample prepared in 25mM Sodium Acetate (pH 5.0),100mM NaCl and dissolved in 90%-10% v/v H_2_O-D_2_O buffer, was concentrated up to 1mM. All NMR spectra were recorded at 298K on 600 MHz Bruker Avance III HD spectrometer equipped with a cryoprobe. The ^15^N-edited HSQC spectrum of vUb showed that the backbone amide resonances were well-dispersed, suggesting folded protein (Fig 2B). A series of 3D NMR experiments including 3D CBCA(CO)NH, HNCACB, HNCA, HN(CO)CA and HNCO were recorded and analyzed to assign chemical shifts of backbone ^1^H, ^15^N, ^13^Cα, ^13^C’ and the side-chain ^13^Cβ resonances. The dihedral angles were obtained by analyzing the backbone chemical shifts in TALOS+ (18). Standard 3D H(CCO)NH, (H)CC(CO)NH and HCCH-TOCSY and HCCH-COSY experiments were carried out to assign the chemical shifts of the side chain atoms. 2D (HB)CB(CGCD)HD experiment and 2D ^13^C aromatic HSQC spectra were used for assigning the chemical shifts of aromatic side-chains. ^15^N-edited 3D NOESY-HSQC and ^13^C-edited 3D NOESY-HSQC experiments with mixing time of 100ms were recorded to obtain the distance constraints. All NMR data were processed in Bruker Topspin3.5pI5 and analyzed by NMRFAM-SPARKY software (19). Following peak-picking of the backbone and side-chain experimental data in SPARKY, the peaks were assigned manually. Automatic peak picking was performed for NOESY spectra using NMRFAM-SPARKY. NOESY peak assignments were performed in CYANA (20) and corrected over multiple cycles of structure building. In the final cycle, 100 structures were calculated using dihedral and distance constraints, and the twenty lowest energy structures superimposed with rmsd 1.1Å (Fig 2C). The backbone dihedral angles of the final converged structures were evaluated by the Molprobity and PSVS suite of programs (21, 22). The NMR and refinement statistics are provided in Table 1.

**Figure 2.**
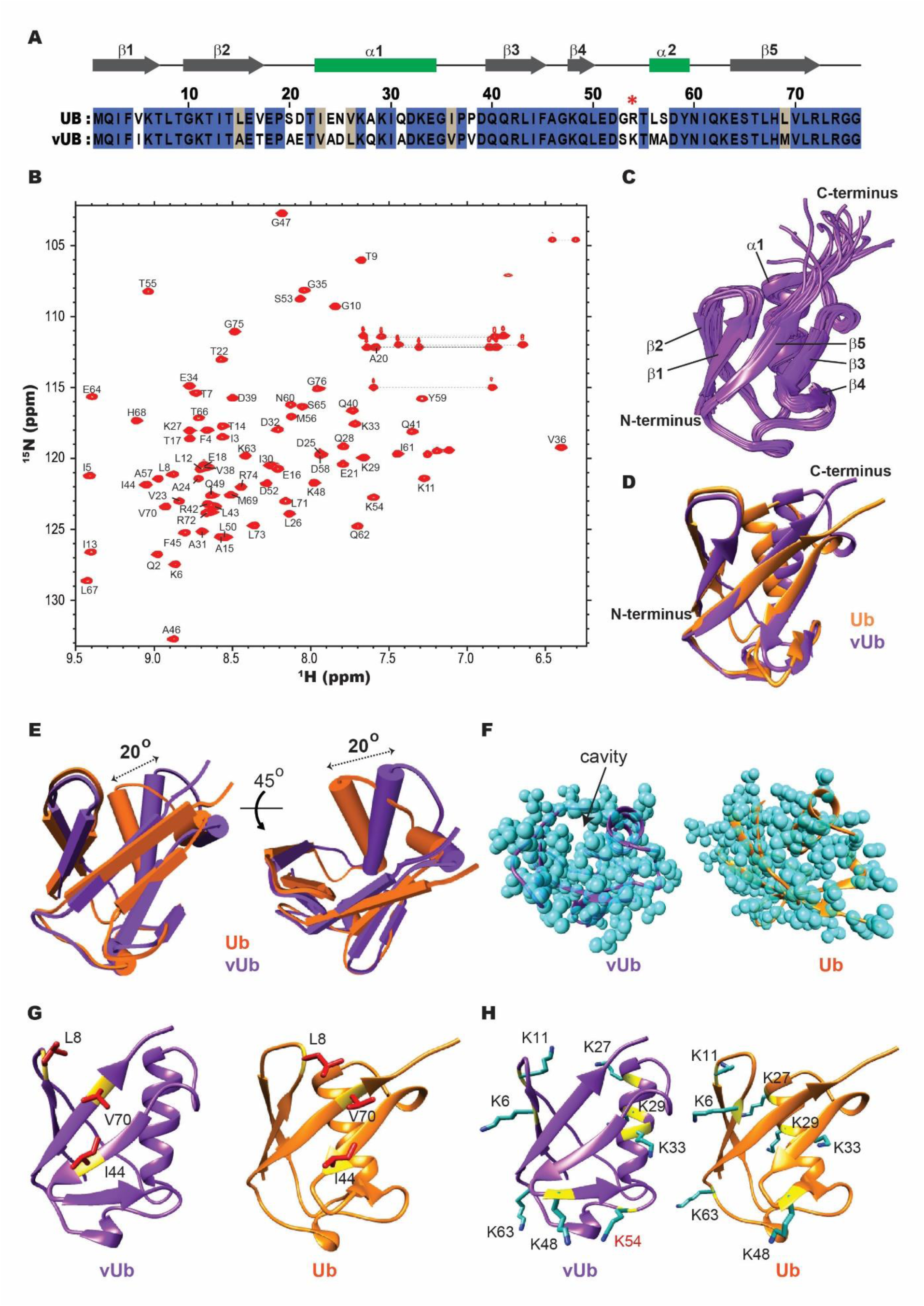
Solution structure of vUb. (A) Alignment of human Ub and vUb with secondary structure information given on the top. The identical residues are colored in blue in the background. The non-conserved hydrophobic residues are colored in gray in the background. (B) The ^15^N-edited HSQC spectrum of vUb is shown. The assigned backbone amide resonances are represented by one-letter amino acid code followed by residue number. The side-chains resonances of glutamines and asparagines are connected with by black dashed lines. (C) The twenty lowest energy structures of vUb are in cartoon representation. The N-termini, C-termini, and secondary structures are labeled. (D) The lowest energy structure of vUb (purple) is superimposed on the structure of Ub (orange). (E) The vUb/Ub superposition is shown in pipes-and-planks representation. There is a difference of twenty degrees in the orientation of helix α1 between Ub and vUb. (F) The structures of Ub and vUb are shown as space-filling models, where the atoms are shown as cyan spheres. (G) The important hydrophobic patch L8-I44-V70 is shown for Ub and vUb. The sidechains of the residues are shown in red. (H) The lysines used to extend polyubiquitin chains in Ub and vUb are shown. The sidechains are colored in blue. The lysines are colored in black, and the additional lysine K54 in vUb is colored in red.

**Table 1.** NMR and refinement statistics of vUb

|  |  |  |
| --- | --- | --- |
| **Distance Restraints (NOE)** |  |  |
| Short range | ($\vert i-j\vert \leq 1$) | 1206 |
| Medium range | ($\vert i-j\vert < 5$) | 136 |
| Long range | ($\vert i-j\vert \geq 5$) | 204 |
| Total |  | 1344 |
| **Other Restraints** |  |  |
| Dihedral Angles ($\Psi, \Phi$) | | 130 |
| CYANA target function ($\text{\AA}^2$) | | 0.27($\pm 0.04$) |
| **Average Pairwise Rmsd<sup>a</sup> ($\text{\AA}$)** |  |  |
| All Backbone atoms |  | 0.6 |
| All Heavy atoms |  | 1.1 |
| **Deviations from idealized geometry** |  |  |
| Bond Angles ($^\circ$) | | 0.2 |
| Bond lengths ($\text{\AA}$) | | 0.001 |
| Close Contacts |  | 0 |
| **Ramachandran Statistics<sup>a</sup>** |  |  |
| Most Favoured regions (%) |  | 97 |
| Allowed regions (%) |  | 3 |
| Disallowed regions (%) |  | 0 |

<sup>a</sup> Calculated for ordered residues (2-72) in an ensemble of 20 lowest energy structures.

The structure of vUB forms the conserved beta-grasp fold like ubiquitin, including five beta sheets (β1-β5) and one helix (α1) (Fig 2C). The structure of vUb and Ub superimposes well with an rmsd of 2.0 Å (Fig 2D). A significant difference in structures of Ub and vUb is observed in the orientation of helix α1 (Fig 2E). Due to the substitutions in β-strands β2 and β5, and helix α1, there are differences in the side-chains present at the buried core of the protein (Fig S1A & S1B). These differences give rise to altered packing, and consequently, the helix α1 in vUb is rotated by twenty degrees away from the helix α1 in Ub (Fig 2E). The helix α1 in Ub is closer to the β1-β5 beta-sheet than α1 in vUb. As a result, a cavity is observed in vUb, and the two salt-bridges present in helix α1 of Ub are absent in vUb (Fig 2F and S1C). The electrostatic surface potential distribution is similar, except a minor additional negative charge is observed in the loop between β1 and α1 of vUb (Fig S1D). The negatively charged residue D21 is buried in Ub, but due to differences in packing of the loop between β1 and α1, the corresponding residue E21 is exposed in vUb giving rise to the minor difference in surface charge. The surface of beta-sheet that presents the critical hydrophobic patch is similar between Ub and vUb (Fig 2G). Apart from the seven conserved lysines, an extra lysine K54 is present in the loop between β4 and α2 in vUb (Fig 2H). Overall, the structural studies suggest that due to the cavity at the buried core and the absence of salt-bridges, the stability of vUb may be different from Ub. On the other hand, since the hydrophobic surface patch in conserved in vUb, the activation of vUb by enzymes in the Ubiquitination reaction and interaction of vUb with Ubiquitin binding domains may be indifferent between vUb and Ub.

### The backbone dynamics of vUb is similar to Ub

The backbone dynamics in the ps-ns timescale of vUb and Ub was compared by measuring the Longitudinal (T1) and transverse (T2) relaxation and the heteronuclear nuclear Overhauser enhancement (het-NOE) of the two proteins. The dynamics experiments were carried out at 298K and pH 6.0 (Fig 3A-3D). These measured values were used to analyze the order parameter (S^2^) using the model-free approach (23, 24). The values of S^2^ are limited between 0 and 1, where S^2^ close to 0 means a higher degree of disorder in N-H bond vector, and value close to 1 indicates restricted N-H bond vector. The average S^2^ value of vUb is 0.824, indicating that the protein is significantly rigid. A minor decrease in the S^2^ values was detected in the loop between β1 and β2, and the loop between β4 and β5, suggesting that these regions are more dynamic than the rest (Fig 3E). For comparison, the T1, T2, and het-NOE experiments were repeated on ^15^N-Ub, and its S^2^ was calculated (Fig S2A-S2E). A comparison between the S^2^ values of Ub with vUb is given in Fig 3F. The regions in vUb that have values higher than standard deviation include the region in between β2 and α1, and the region from β4 to β5, indicating that these regions are more dynamic in vUb than Ub (Fig 3F). The lower rigidity in the loops is in synch with the differences in packing observed in the structure of vUb. Interestingly, the loop between β3 and β5 is dynamic and disordered in a near-native folding intermediate of Ub, which has lower stability than native Ub (25). Hence, the higher dynamics in this region of vUb suggests that Ub may be more stable than vUb.

**Figure 3.**
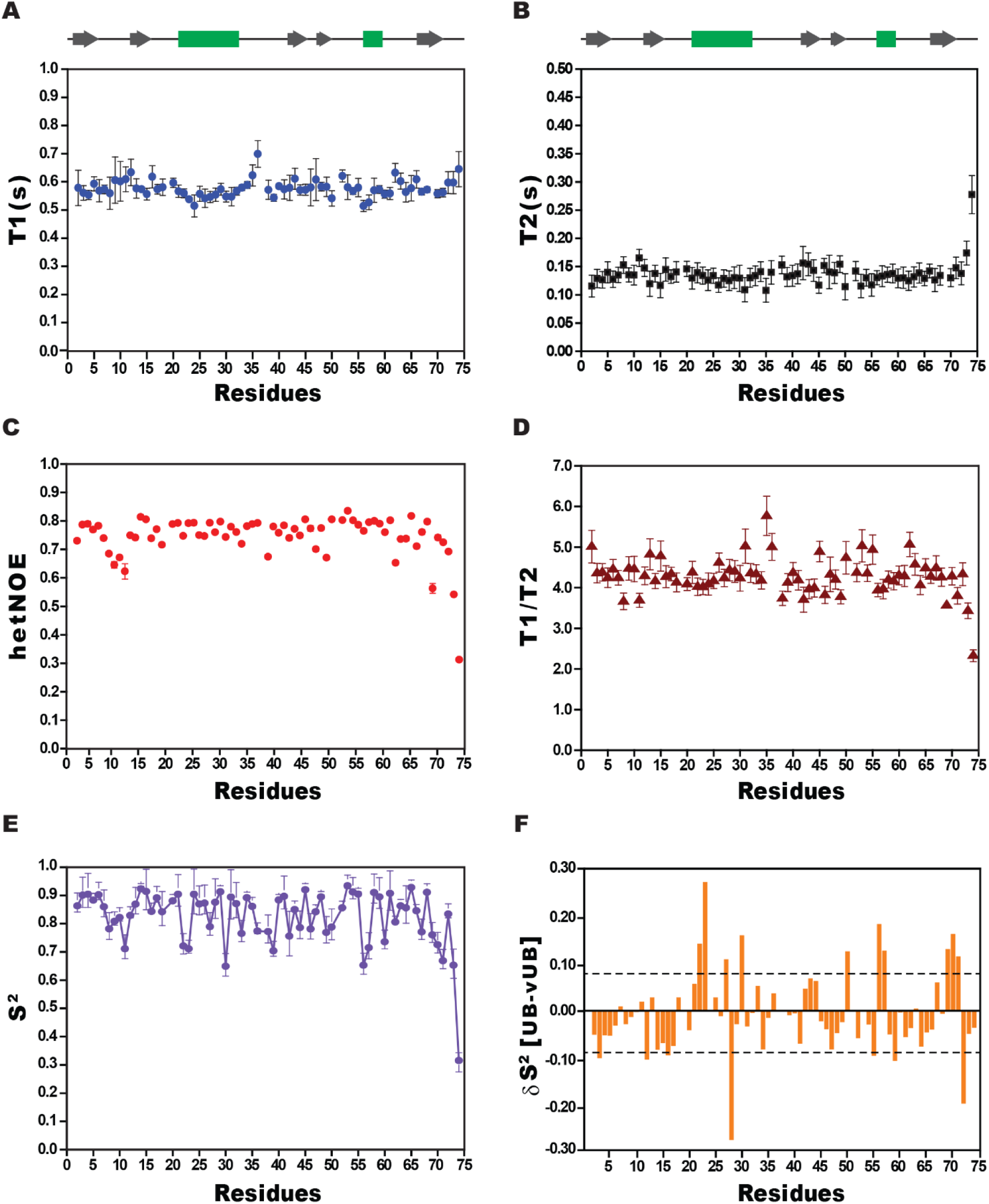
The ps-ns backbone dynamics of vUb. The measured values of (A) longitudinal relaxation, T1, (B) transverse relaxation, T2, and (C) heteronuclear nuclear Overhauser enhancement, het-NOEs, were plotted for each residue of vUb. (D) The T1/T2 ratio for each residue is plotted against the residue number of vUB. (E) The order parameter S^2^ was calculated using the Lipari Szabo model and plotted against the residue number of vUB. (F) The difference in S^2^ between Ub and vUb (δS^2^) is plotted. The standard deviation is highlighted with the black dashed line. The secondary structure elements of vUb against its sequence are provided on top of each plot.

### Ub is more stable than vUb

Chemical-induced and temperature-induced denaturation melts were carried out to compare the stability of vUb and Ub. For the chemical denaturation melt, purified vUb (40 µM) and Ub (40 µM) were incubated overnight in different concentrations of guanidium chloride (ranging from 0-6 M). Subsequently, Circular Dichroism (CD) spectra were collected for the proteins. The concentration of Guanidium chloride required to unfold 50% of vUb was 1.9 M. Alternately, Ub required 3.6 M Guanidium chloride to unfold 50% of the protein, indicating that Ub was more stable than vUb. Hence, ΔG° reduced from 11.5 kcal mol^-1^ for Ub to 6.6 kcal mol^-1^ for (Fig 4A and Table 2). The stability of the proteins was also measured by thermal denaturation. The melting point (Tm) for vUB was 74°C. Ub did not melt up to 90°C, confirming it is more stable than vUb (Fig 4B).

**Figure 4.**
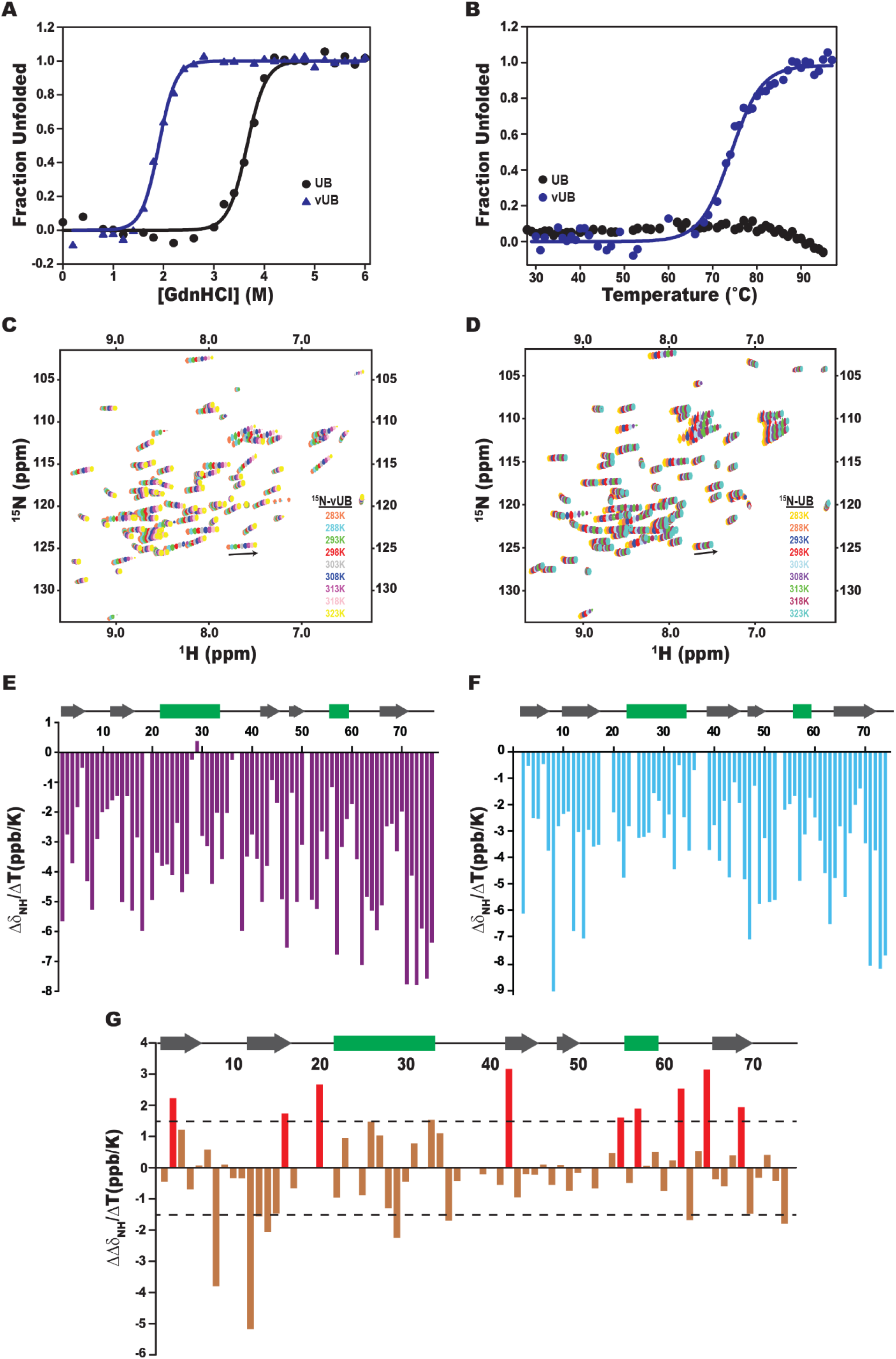
A comparison of the stability of Ub and vUb. (A) GdnCl melt curves of vUb and Ub. Normalized mean ellipticity shift is plotted against GdnCl concentration (B) The thermal melt curve of vUb and Ub. Change in ellipticity is normalized and plotted against temperature. (C) & Overlay of ^15^N-HSQC of vUb and Ub, respectively, at temperatures ranging from 283K to 323K. Temperature coefficients are plotted against the residue numbers for (E) vUb and (F) Ub. The secondary structure elements of vUb and Ub are drawn on top of the respective plots. (G) The difference in temperature coefficients of Ub and vUb is plotted against residue numbers. The standard deviation is shown as dashed black lines. Red bars represent the residues in vUb that are significantly destabilized upon increasing temperature. The secondary structure elements of vUb are drawn on top of the plot.

**Table 2.** Analysis of Guanidium chloride melting curves of Ub and vUb

| | $\Delta G_0$ (kcal mol<sup>-1</sup>) | m (kcal mol<sup>-1</sup>) | [GdnCl] (in M) at 50% unfolding |
| --- | --- | --- | --- |
| Ub | 11.5±1.2 | 3.2±0.3 | 3.65 |
| vUb | 6.6±0.5 | 3.4±0.2 | 1.89 |

The local stability of vUb and Ub were probed using NMR by measuring and comparing the temperature coefficients per residue. The 2D-HSQC experiment was recorded for ^15^N-vUb and ^15^N-Ub from 283K till 323K at every 5K increment (Fig 4C and 4D). The resonance shifts were fit linearly with respect to the temperatures to calculate the temperature coefficients (Fig 4E and Fig 4F). The average temperature coefficient ΔδNH/ΔT values of almost all the secondary structures of vUb and Ub were nearly the same. The ΔΔδNH/ΔT (Ub-vUb) values are plotted in Fig 4G, where negative and positive values indicate residues in vUb with higher and lower stability than Ub residues. Intriguingly, the of ΔΔδNH/ΔT values of non-conserved residues like V23, L26, and A31 in α1, T55, and M57 in α2, Q62, S65 and M69 in β5 strand were positive, indicating that these substitutions cause local instability in vUb (Fig 4G). Several residues had high positive values in α2, β5, and in the loop in between, implicating vUb is relatively unstable in these loop regions (Fig 4G).

As mentioned above, the difference in stability between vUb and Ub could be due to the difference in hydrophobic core packing. Alanine scanning studies on Ubiquitin has suggested that hydrophobic contacts at the buried core are essential to maintain high stability and rigidity (26). Mutations of buried residues I30A-Ub and L43A-Ub created a cavity and significantly reduced the stability of Ub. I30 on α1 tightly contacts L15 on β2 strand in Ub (Fig S1A). However, L15 is substituted to A15 in vUb, which disrupts the contact. The I36 residue present in the C-terminal loop of α1 packs against β5 strand by making contacts between its side-chain and the sidechains of L69 and L71 of β5 strand (Fig S1D). L69 is substituted to M69 in vUb, which may loosen contacts between the β5 strand and α1. The higher temperature coefficients of E16 and M69 confirm local instability in these regions due to the substitutions. Altogether, Ub is more stable than vUb. Measurement of backbone dynamics and local stability indicates that the region between β2 and α1, and the region between α2 and β5 are largely dynamic and unstable in vUb when compared to Ub. Often stability of a protein determines its molecular recognition by co-factors (27). The interaction of vUb with Ubiquitin binding domains in proteasome receptors was examined to assess if the differences in stability are reflected in the interactions with co-factors.

### vUb interacts with proteasome receptors Rpn10/S5a and Rpn13

The Ubiquitin-26S proteasome pathway targets misfolded or tightly regulated proteins for degradation by conjugating them with a poly-Ub chain (28). The ubiquitinated substrate interacts with non-ATPase subunits such as Rpn1, Rpn10/S5a, and Rpn13 of the proteasome *via* Ub. Consequently, the substrate enters the core particle for degradation, and the Ub chains are cleaved off by Deubiquitinases (DUBs) present in the proteasome (29–31). Since the insect proteasome subunits have a high degree of similarity to the mammalian system (32), and the proteins of mammalian proteasome receptors were available, the interaction between vUb and the ubiquitin-binding domains in proteasome receptors (S5a and Rpn13) were tested to examine if substrates conjugated with poly-vUb chains can be recognized by the proteasome receptors.

NMR spectroscopy and Isothermal Titration Calorimetry (ITC) was used to determine if vUb binds to Rpn10/S5a. Uniformly labeled ^15^N-vUb was titrated with unlabeled S5a containing the two Ubiquitin Interacting Motifs (UIMs, UIM1: 206-230aa; UIM2: 274-300aa), and 2D-HSQC experiments were recorded till saturation. The change in the chemical environment of vUb residues upon binding with S5a altered the chemical shifts on the backbone amide resonances (Fig 5A). The difference in chemical shifts between the apo and bound form of vUb was plotted as chemical shift perturbation (CSP) in Fig 5B. The hydrophobic surface patches of vUb consisting I44, V70, and L71 along with polar residues of β4 (K48 and Q49) had maximum CSPs, suggesting that these residues are present at the interface of S5a/vUb complex (Fig 5C). Additionally, the hydrophobic residues like A15, I30, L43, G47 and L67 and charged residues like D32, R42, and R72 are also present at the interface. The chemical shifts values were fit against the ligand concentration in the NMR titration experiments to yield the dissociation constant (K_d_) of 78±12 µM.

**Figure 5.**
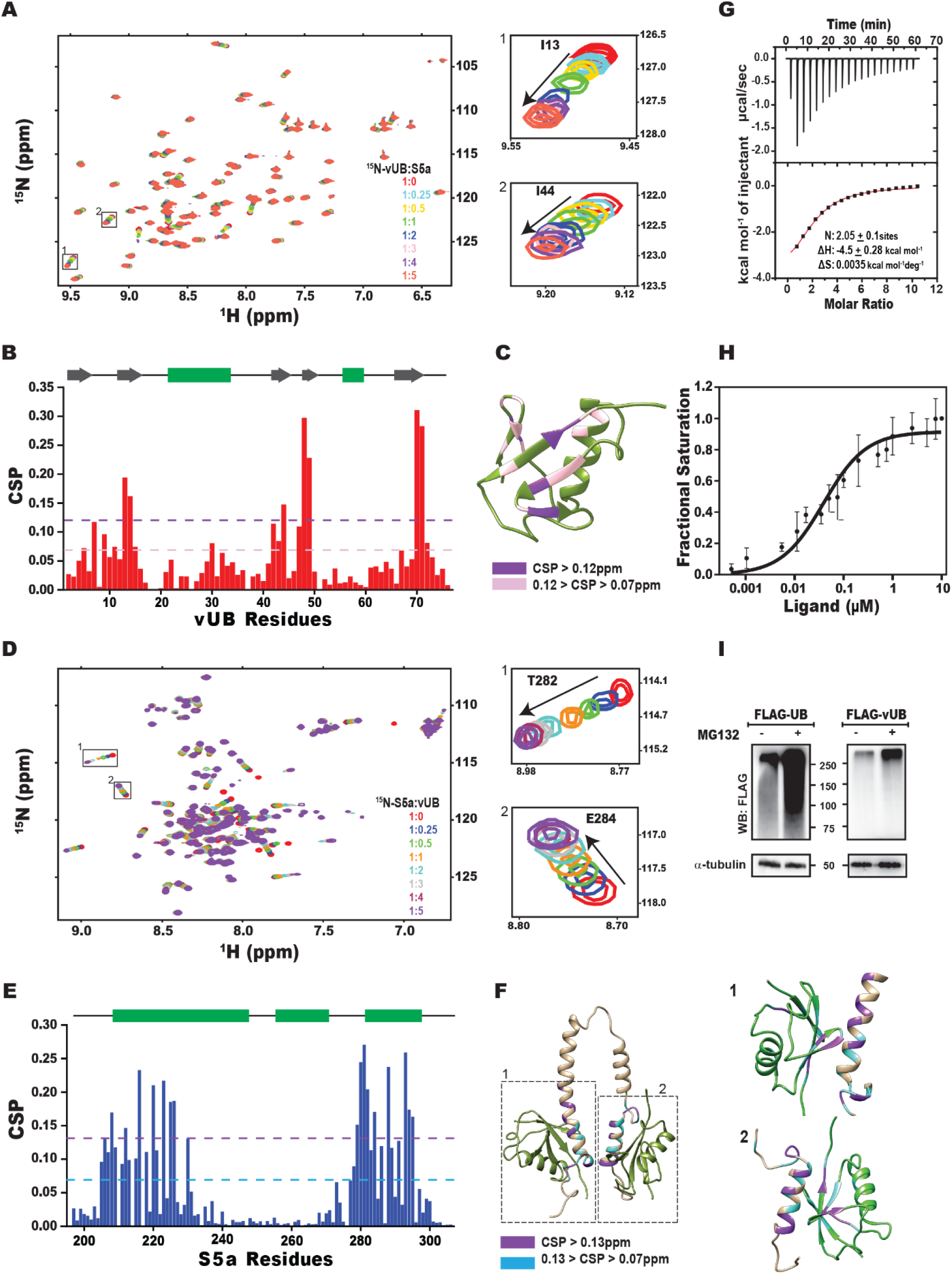
vUb binds to proteasome receptor S5a and Rpn13. (A) Overlay of ^15^N-edited HSQC spectra of ^15^N-vUb with different stoichiometric ratios of S5a. The resonances of residues I13 and I44 are zoomed for clarity. (B) The CSP observed in vUb upon titration with S5a is plotted against vUb residue number. Mean and mean with standard deviation are highlighted in pink and purple, respectively. (C) The high CSPs are mapped on to the vUb structure. (D) Overlay of ^15^N-HSQC spectra of ^15^N-S5a with different stoichiometric ratios of vUb. T282 and E284 resonances are zoomed for clarity. (E) The CSP observed in S5a upon titration with vUb is plotted against S5a residue number. Mean and mean with standard deviation are highlighted in blue and purple, respectively. (F) The residues with high CSPs are highlighted in the S5a structure. Best model structure of vUB (olive green) superimposed on proximal and distal UB in S5a-K48 linked-diUb NMR structure (PDB: 2KDE). Zoomed images show the interaction of helix α1(1) and α3(2) of S5a with vUb. (G)The interaction between vUb and S5a is measured by ITC. The titration curve indicates a 1:2 stoichiometric of S5a:vUb complex. The fit of the ITC data yielded the dissociation constant (K_d_) = 85 (± 9.9) μM, stoichiometry n= 2.0 (± 0.1), ΔH = −4.5 (± 0.3) kcal mol^-1^ and TΔS= 0.09 kcal mol^-1^. (H) The binding curve for Rpn13 Pru domain binding to vUB as determined by intrinsic tryptophan fluorescence, which yielded the dissociation constant of 40 (±14) nM. (I) Accumulation of Ub and vUb conjugates in HEK293T cells upon proteasomal complex inhibition. Cells were transfected with FLAG-Ub or FLAG-vUb, and post inhibition by the proteasomal inhibitor MG132, cell lysates were separated on SDS-page and immunoblotted with anti-FLAG antibody. α-tubulin was used for loading control.

In the reverse titration, uniformly labeled ^15^N-S5a-UIMs (196-306aa) was titrated with unlabeled vUb (Fig 5D). High CSPs were observed in both the UIMs (UIM1: 206-230 aa; UIM2: 274-300 aa), suggesting that both the UIMs interacted with vUb (Fig 5E). The fit of chemical shifts against ligand concentration yielded different K_d_ values, 331±25 µM for UIM1 and 21±6 µM for UIM2. Clearly, UIM2 interacts with vUb with higher affinity. A similar mode of interaction was also observed in S5a/Ub complex, where UIM2 interacted tightly to S5a than UIM1 (31, 33). Residues in the conserved hydrophobic patch (L^216^ALAL^220^) have high CSPs in UIM1 and interacts with vUb. Additionally, several polar and charged residues in UIM1 (R221, S223, E225, and E226) also interact with vUb. Similarly, apart from the hydrophobic patch (I^287^AYAM^291^) in UIM2, a series of polar and charged residues (S279, S280, E284, E285, Q292, S294, Q296, and E299) are also involved in the interaction with vUb (Fig 5F). Overall, the affinity and the interface of the vUb/S5a complex is similar to the Ub/S5a complex, indicating that S5a can identify substrates conjugated with vUb. The binding of vUb and S5a-UIMs was further confirmed by ITC (Fig 5G). The binding isotherm was fitted to a 1:2 complex to obtain the overall K_d_ of 85.0±9.9 µM, which agrees well with the values as calculated by ^15^N-vUb/S5a NMR titration. The ΔG values were calculated to be −4.6 kcal mol^-1^.

The binding interaction of vUb with another proteasome receptor, Rpn13 was detected by NMR. The N-terminal domain of hRpn13 called Pru domain (1-150 aa) interacts with Ub (30). Uniformly labeled ^15^N-vUb was serially titrated with hRpn13-Pru domain from 1:0.5 till 1:4 molar ratios and their subsequent 2D ^15^N-edited HSQC spectra were recorded (Fig S3A) and the CSP plot is shown in supplementary fig S3B. Amide resonances of residues L8, T14, R42, K48, Q49, and L71, disappeared during titration suggesting peak broadening due to conformational exchange. Resonances of F4, F45, A46, H68, and V70, disappeared during the initial titration points, but reappeared in the later titration molar ratios, suggesting the intermediate exchange and high affinity of the interaction. A similar observation was made in the NMR titration of Rpn13/Ub complex (30).The dissociation constant of Rpn13/vUb complex was measured by fluorescence spectroscopy, where post excitation at 295 nm, the tryptophan spectra of Rpn13 was monitored from 300 nm to 400 nm. The fluorescence-based titration yielded the K_d_ 40+15 nM, (Fig 5H) which is similar to the affinity between Rpn13 and Ub. The interaction of vUb with proteasome receptors like S5a and Rpn13 is indifferent from Ub (Table 3), which suggests that proteins conjugated with poly-vUb chains can be identified by the proteasome.

**Table 3.**
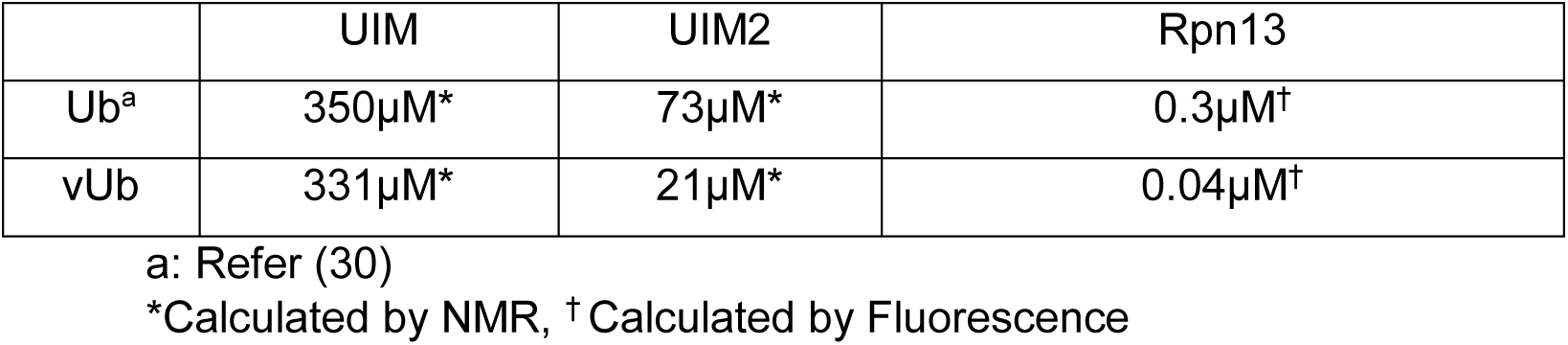
Ub binding with Rpn10 and Rpn13

The degradation of vUb conjugated substrates in cellular conditions was studied by transfection of HEK293T cells with FLAG-Ub and FLAG-vUb, followed by detection of the polyUb and poly-vUb conjugated substrates in the presence/absence of proteasome inhibitor MG132. Similar to the Ub-conjugated substrates, the vUb conjugated substrate accumulate significantly upon addition of proteasome inhibitor, indicating that these conjugates are recognized and degraded by the proteasome (Fig 5I). These results indicate that vUb can bind to Ubiquitin binding domains in the proteasome receptors, and can act as a signal for proteasomal degradation.

### Polyubiquitination activity of vUb is similar to Ub

It is important to investigate if the differences in structure and stability can affect the activation of vUb by enzymes in the Ubiquitin pathway. The poly-ubiquitination activity of vUb was measured using multiple E2:E3 pairs. Initially, E2s that form specific Ubiquitin linkages were used in the ubiquitination reactions. E2-25K specifically forms K48-linked poly-Ub chains whereas, Ubc13 (along with its co-factor Mms2) specifically forms K63 linked poly-Ub chains. E2-25K and Ubc13 can synthesize poly-Ub chains without any E3 *in-vitro* (34, 35). Ubiquitination reactions were carried out with Ube1, E2-25K, and either Ub or vUb in the reaction mix. The intensity of polyubiquitin chains observed in the western blots was similar between Ub and vUb, indicating that vUb forms K48-linked chains with equal efficiency as Ub (Fig 6A). The amount of K63-linked Ubiquitin chains catalyzed by the Ubc13/Mms2 complex was less in vUb than Ub (Fig 6B).

**Figure 6.**
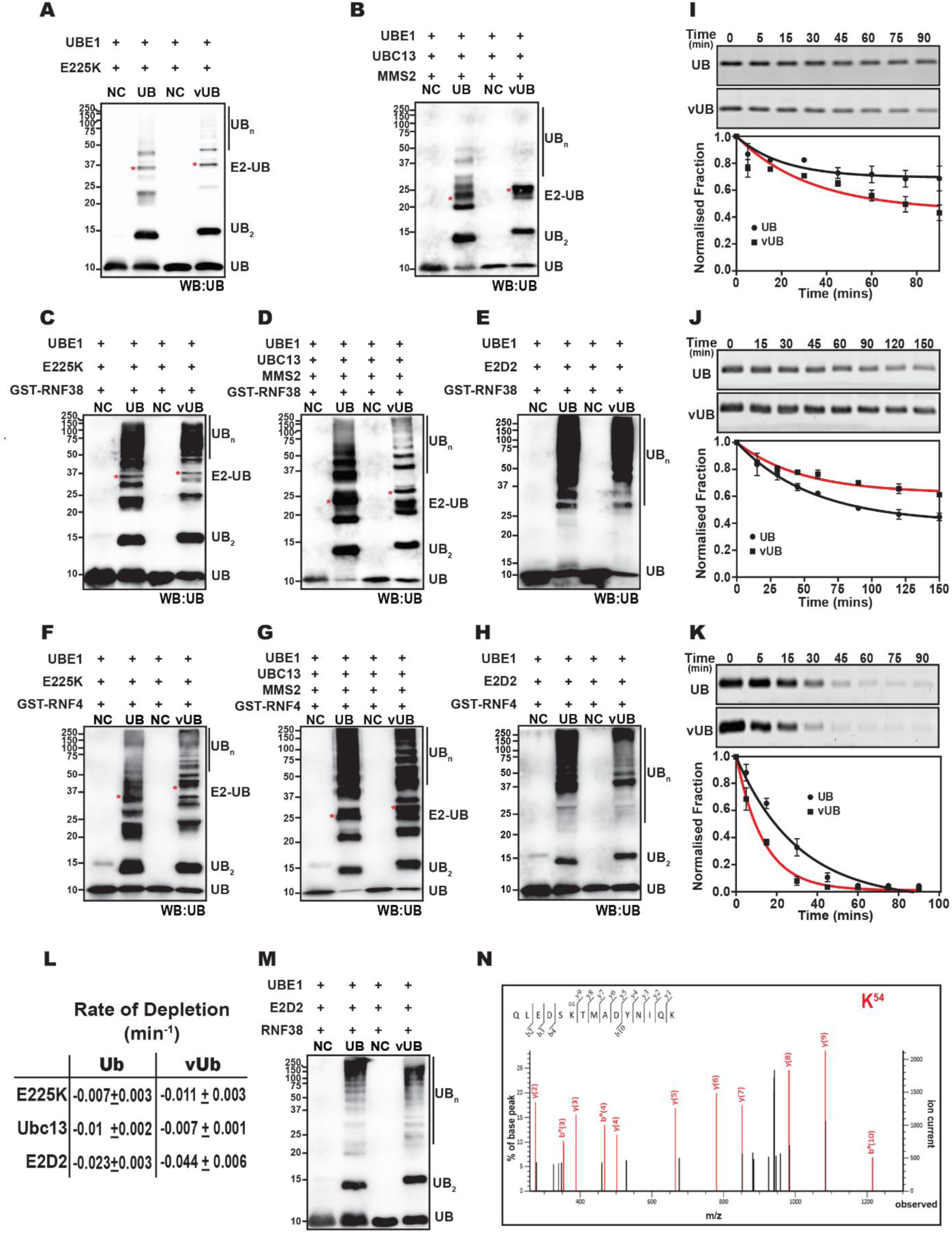
A comparison of the polyubiquitination activity between vUb and Ub. The polyubiquitination reaction with either Ub or vUb by (A) E2-25K, (B) Ubc13/ Mms2, (C) E2-25K/RNF38, (D) Ubc13/ Mms2/RNF38 and (E) E2D2/RNF38. (F), (G) and (H) similar as (C), (D) and (E), respectively, where RNF38 is substituted by RNF4. All the reactions were probed with the anti-Ub antibody. (I) Depleting amount of mono-Ub (top) and mono-vUb (down) against time were detected in a polyubiquitination assay using E2-25K and RNF38 as the E2 and E3, respectively. The amount of mono-Ub and mono-vUb are plotted at the bottom. (J) & (K) is the same as (I), where the E2 is Ubc13/ Mms2 and E2D2, respectively. (L) The rates of depletion of UB and vUB have been calculated by fitting the initial time points, whose slope represents the rates. (M) Confirmation of di-vUb formation using E2D2 and RNF38 by immuno-detection with the anti-Ub antibody. The same reaction was separated on SDS page gel, and the di-vUb was excised for MS/MS analysis. (N) MS-MS spectrum of one of the tryptic peptide (Seq: QLEDS**K**^54^TMADYNIQK, MW: 1896.8887Da, 2+) that confirms the isopeptide formation by K54^th^ residue.

E3s catalyze the rate of ubiquitination by several folds. The *in-vitro* reactions were repeated in the presence of two different E3s, RNF38, and RNF4. The synthesis of K48-linked polyubiquitin chains was indifferent between vUb and Ub when either RNF4 or RNF38 is used (Fig 6C and Fig 6F). The synthesis of K63-linked vUb chains was, however, slightly lower in the presence of E3s (Fig 6D and Fig 6G). Finally, E2D2 was used as the E2 in the ubiquitination reaction. E2D2 synthesizes poly-Ub chains of various linkages. Polyubiquitination by E2D2/RNF38 or E2D2/RNF4 complexes were similar between Ub and vUb (Fig 6E and Fig 6H).

Analysis of the polyubiquitin chains may fail to detect minor differences in the activation of Ub and vUb. As mono-Ub is activated by the enzymes to form poly-Ub chains in ubiquitination reaction, the rate of mono-Ub depletion reflects the efficiency of activation by Ubiquitin enzymes. Hence, the depleting mono-vUb or monoUb species was quantified over time in a typical polyubiquitination reaction. These experiments were performed with the E2s E2-25K, E2D2, and Ubc13/Mms2, using RNF38 as the E3. Interestingly, minor differences were observed in the activation of Ub and vUb, depending on the E2 used in the reaction. The activation of vUb is slightly better than Ub in the case of E2D2 and E2-25K (Fig 6I and 6K). However, in the presence of Ubc13/Mms2, activation of vUb was slower than Ub (Fig 6J). In all the assays, the activity of Ubc13/Mms2 complex was lower for vUb than Ub. Unlike the E2-25K and E2D2, the activity of Ubc13/Mms2 depends on the interaction between Mms2 and acceptor Ub, which could be different between Ub and vUb. Overall, the rate of depletion in Ub did not differ by more than 2-fold over vUb, indicating that the activity of Ub was similar to vUb (Fig 6L).

The activation of Ub by E1 and the trans-thiolation to E2 depends on the hydrophobic patch and the C-terminal end residues (36). Since these are conserved in vUb, the E1 activation and transthiolation activity is expected to be similar between Ub and vUb. R54 in Ub contributes to the intrinsic affinity of Ub with E1. Replacing this charged residue with Leucine affects the binding of E1 with Ub (37). However, if replaced with a charged residue like lysine, the binding affinity is not affected (38). Hence, although vUb has lysine at 54^th^ position, and conjugation capability with E1 and E2 should be similar. Finally, the activation of E2∼Ub for transfer of Ub to substrates is induced by the closed conformation of E2∼Ub, where the central helix in the UBC domain of E2 contacts the hydrophobic patch of Ub. Since the hydrophobic patch is conserved in structure, it is reasonable that the ubiquitination activity of vUb is similar to that of Ub.

### Polyubiquitin chains formed by vUb includes a novel K54-linked chain

The substitution of R54 by K54 in vUb provides an interesting possibility, where a new linkage type (K54-linked) polyubiquitin chains are formed in vUb. E2D2 can synthesize all the varieties of linkage chains K11, K48, K63. The poly-vUb chains catalyzed by E2D2 may include K54 linked vUb chain. A reaction was carried out using E2D2, RNF38, and vUb. After the reaction, the products were separated on SDS page gel and simultaneously blotted with Ub antibody (Fig 6M). The band corresponding to di-vUb was extracted from the SDS gel, digested by trypsin and analyzed by MS/MS. The MS/MS spectra detected multiple types of isopeptide linkages *viz.* K6-, K11-, K48-, and K63- (Table 4 and Fig S4A-S4D). Apart from these known isopeptide linkages, a peptide containing the novel K54-linkage in di-vUb was also detected (Fig 6N).

**Table 4.**
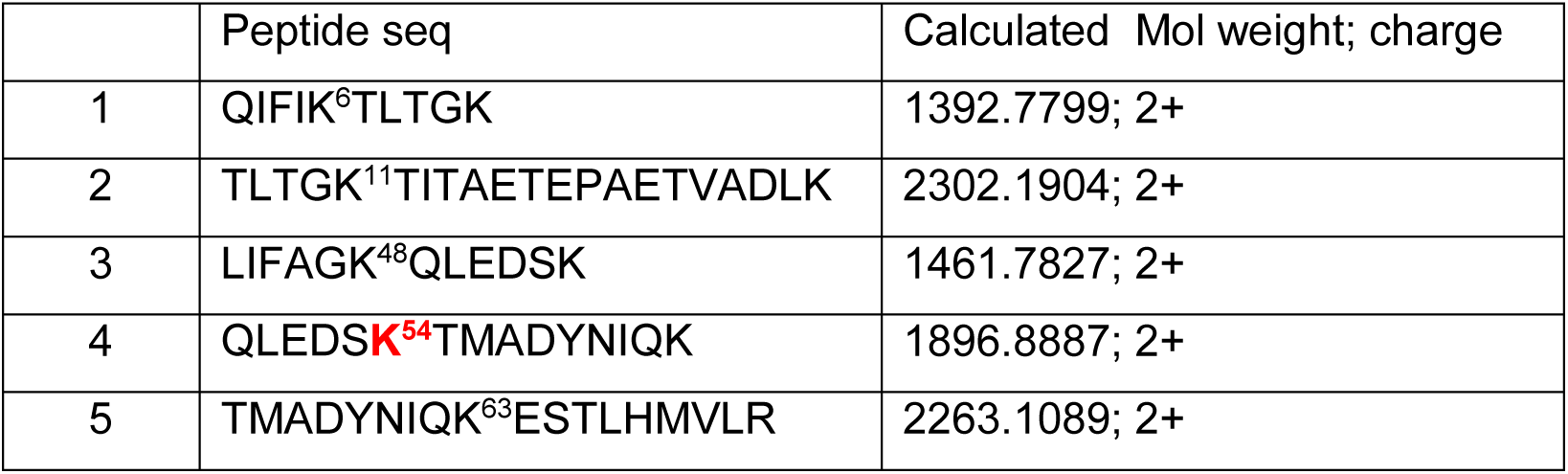
List of peptides of di-viralUb

To further confirm the K54-linked chains, a mutant was designed in vUb, where all the lysines except K54 were substituted to arginines (K54only-vUb). The far UV-CD scan of the K54only-vUb protein confirmed that secondary structure is retained (Fig S5A). Structurally, K54 lies in between K48 and K63. Given that K54-linked chains are unique, whether the E2s that build specific type chains can assemble K54-type chains is unknown. The ubiquitination reactions with E2-25K or Ubc13/Mms2 were repeated with K54only-vUb. Intriguingly, K54-linked di- or poly-vUb were not detected in this reaction (Fig 7A and 7B). The K54-linked chains were not detected even when the E3s RNF4 was used (Fig 7C and 7D). Hence, the E2s that catalyze specific Ubiquitin linkages cannot catalyze K54-linked chains, indicating that these chains may have a distinct topology from K48- and K63-linked chains. Being promiscuous in catalyzing a variety of linkages, E2D2 could catalyze K54-linked poly-vUb chains (Fig 7E, S5C). Apart from K54 linked, these chains could also be N-terminal linked. However, no polyubiquitin chains were observed when K0-vUb was used in the ubiquitination reaction, indicating that the chains in Fig 7E are linked by K54 (Fig 7F). Moreover, the K54-linked di-vUb was analyzed by MS/MS, which confirmed the isopeptide linkage at K54 (Fig 7G). Altogether, vUb can form an entirely new K54-linked polyubiquitin chain, whose topology could be different from the previously known linkages.

**Figure 7.**
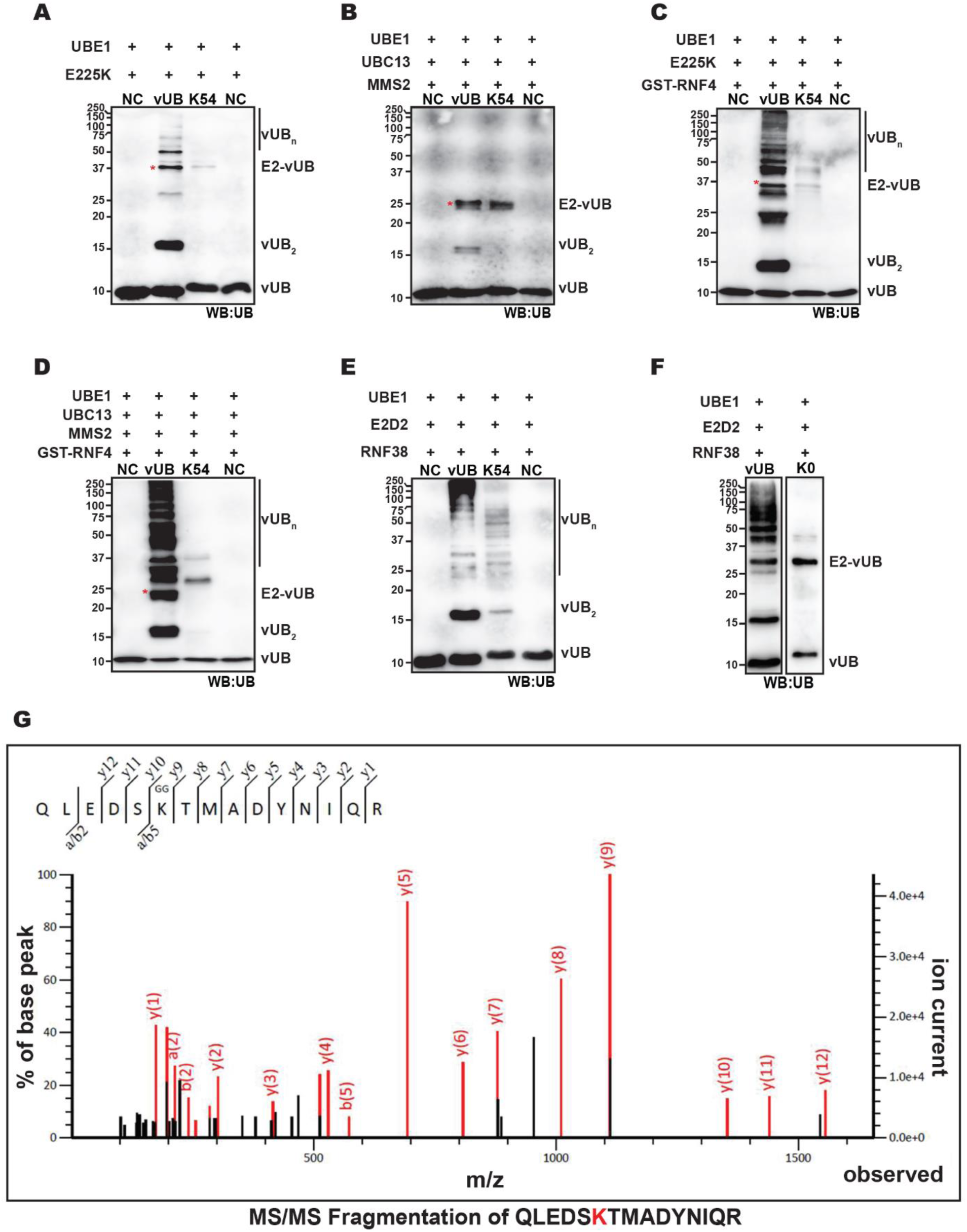
Comparison of poly-ubiquitin conjugate formation between vUb and K54-vUb mutant. Poly-ubiquitination reaction with either wt-vUb or K54-vUb mutant by (A) E2-25K and (B) Ubc13/Mms2. (C) and (D) same as (A) and (B), respectively, but in the presence of E3 RNF4. Poly-vUb conjugate formation by K54-vUb mutant with E2D2/RNF38. The reaction was separated on an SDS page gel, and the di-vUb band was excised for MS/MS analysis. (F) Reaction same as E but K0-vUB mutant is used in the second lane. (G) MS/MS spectrum of a tryptic peptide (Seq: QLEDS**K**^54^TMADYNIQR, MW: 1924.8949Da, 2+) confirmed K54-linked di- vUb in the reaction.

### K54-linked Ubiquitin chains are not cleaved by DUBs

Since K54-linked Ubiquitin chains are not formed by the eukaryotic Ub, they may not be susceptible to cleavage by the host DeUbiquitinases (DUBs). The poly-vUb chains and K54-linked poly-vUb chains formed by E2D2 and RNF38 were treated with three different DUBs (USP2_CD_, AMSH, and OTUBAIN). USP2_CD_ is a non-specific DUB, AMSH is specific for K63-linked chains, and OTUBAIN is specific for K48-chains. USP2_CD_ could efficiently cleave poly-vUb chains, but not K54-linked poly-vUb chains (Fig 8A). The treatment with K63-linkage specific DUB, AMSH, reduced poly-vUb chains partially. This is expected given that AMSH will specifically cleave the K63-linked chains, leaving the other linkages intact. However, the K54 poly-vUb chains were not cleaved (Fig 8B). The similar result was observed when the K48-linkage specific DUB OTUBAIN1 was used (Fig 8C). Overall, the results indicate that the K54-linked poly-vUb chains are fairly resistant to DUBs.

**Figure 8.**
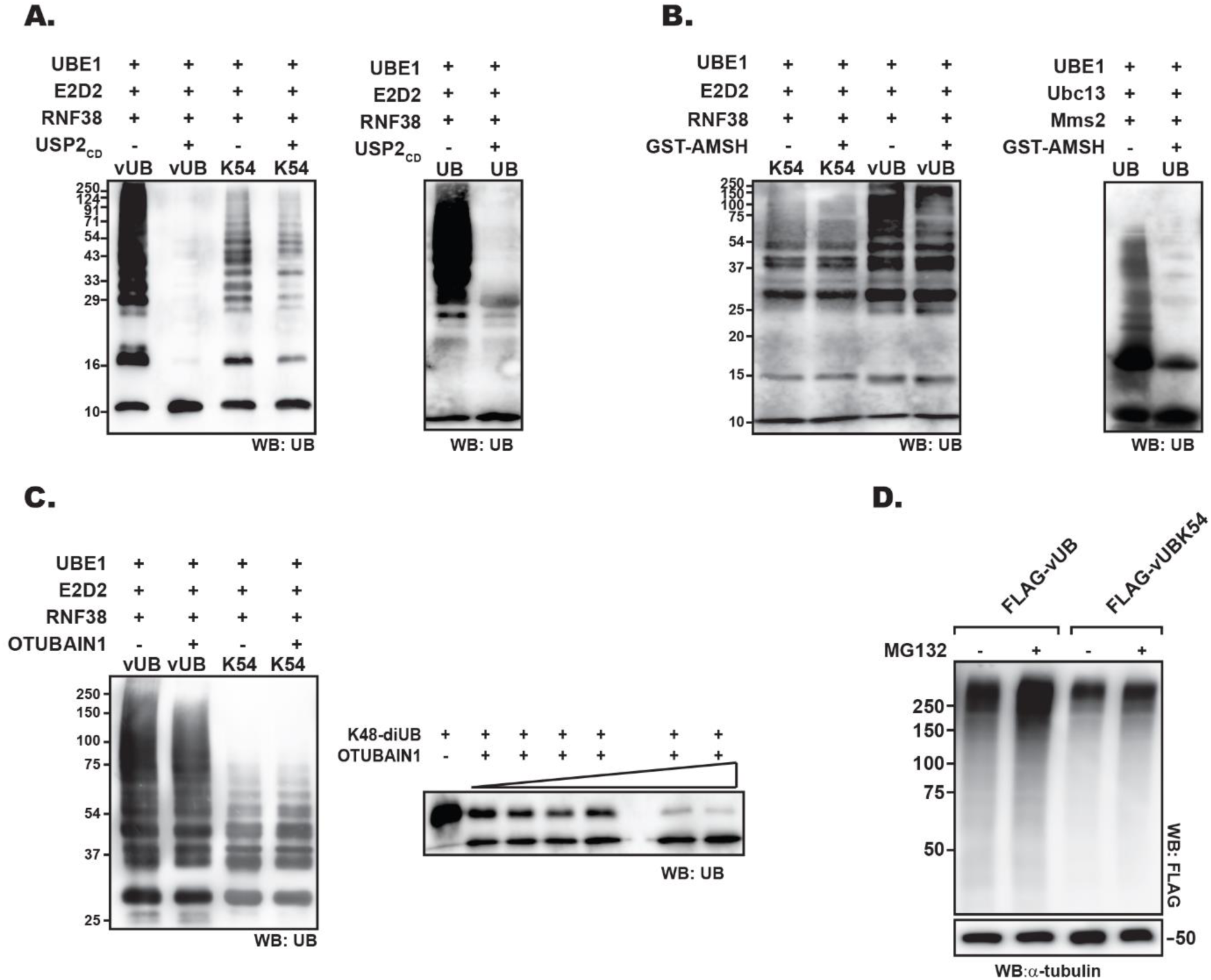
Deubiquitination of vUb conjugates by DUBs. vUb conjugates were formed by ubiquitination reaction involving Ube1, E2D2, and RNF38 and quenched by EDTA. Deubiquitination reaction of these conjugates was carried out using (A) USP2_CD_, (B) GST-AMSH, and (C) OTUBAIN1. The Ub controls for the respective DUBs have been shown next to each image. (D) Accumulation of vUb and K54-vUb mutant conjugates in HEK293T cells upon proteasomal inhibition by MG132. The blots were probed with anti-FLAG antibody to detect FLAG-vUb and FLAG-vUbK54. α-tubulin was used as a loading control.

Whether K54only-polyvUb chains participate in the Ubiquitin-proteasomal pathway was tested by transfecting HEK293T with FLAG-vUb and FLAG-vUb-K54. Cells were subsequently treated with proteasome inhibitor MG132 to detect the accumulated proteasomal substrates. High molecular weight K54only-polyvUb conjugates could be observed in the cells (Fig 8D). However, the amount of K54only-polyvUb conjugates did not differ in the presence/absence of MG132, suggesting that these conjugates may not be targets for proteasomal degradation (Fig 8D). Hence, we uncovered atypical linkage in vUb, that is neither effectively cleaved by DUBs, nor are they targeted by the proteasomal pathway. Conjugation of viral proteins by the vUb is important for assembly of virions (15, 16). Ubiquitination of nucleocapsid proteins specifically with vUb determines whether they will egress from the nucleus to form budded virus or not(17). Preservation of such signals from the host’s antiviral response is essential for the virus. The host can regulate the vUb conjugates by DUBs. Alternately, the host can degrade the vUb conjugated substrates by the Ubiquitin-proteasome pathway. The K54-linked vUb chains may be a counter mechanism of the virus against such antiviral responses.

## Conclusion

Nucleopolyhedrosis viruses (NPV) and *Granulovirus* (GV) belonging to *Baculoviridae* family of viruses encode Ubiquitin in their genomes. Ubiquitin is the central component of the Ubiquitin signaling pathway. Evolutionary analysis of viral Ubiquitin and eukaryotic Ubiquitin from different viruses and host insects have suggested the possibility of horizontal gene transfer, and the phylogenetic analysis further confirms the likelihood of one common ancient ancestral gene (39). However, the retention of vUb indicates that it holds unique properties different from the eukaryotic Ub. We report that differences in the sequence between vUb and Ub have resulted in minor structural differences at the buried core of vUb. Consequently, the stability of vUb is reduced compared to Ub. However, the surface properties of Ub is retained in vUb, and its polyubiquitination activity and interaction with co-factors are indifferent from Ub. Additionally, vUb can form a unique K54-linked chain that is not regulated by either the host DUBs or the host proteasomal pathway. Hence, modification of host/viral proteins by the K54-linked viral Ubiquitin may be an effective way to counter antiviral responses that regulate the viral protein. The molecular details of K54-linked chain, how it resists proteasomal degradation and DUBs, and the function of these chains, are now exciting questions for future research.

## Methods

### Reconstruction of vUb gene tree

39 Baculoviruses containing vUb encoding gene were selected for tree reconstruction, and only the amino acid sequences for Ubiquitin-like protein were selected, whereas the non-ubiquitin amino acid sequences were discarded. The gene tree was reconstructed using the Maximum Likelihood (ML) method. Sequences were aligned using muscle algorithm provided by MEGA X (40). In order to reconstruct the ML phylogeny, we used Partition finder (41) to obtain the evolutionary model for codon evolution (GTR+I+γ, GTR+γ, and HKY+I+γ for three codon positions, respectively). Additionally, RaxML Blackbox (42) was used in the CIPRES platform (43) to get the ML tree.

### Cloning

The DNA sequence encoding vUb and its K54-vUb mutant (in which all lysine were substituted to Arginine, except K54) were synthesized commercially and then cloned into pET3a vector without any tag using NdeI and BamHI restriction sites. The ligation reaction was carried out by T4 DNA ligase, and the plasmid, containing the insert, from the ligated colonies were further verified by Sanger sequencing method using T7 promoter and terminator primers.

### Purification of vUb and vUb-K54

Plasmid encoding vUb and K54-vUb mutant were transformed into BL21 DE3* *E. coli* expression strain. Cells were grown at 37°C initially, followed by IPTG induction and incubated further at 37°C for 5-6 hrs. Harvested cell pellet of vUb was dissolved in Sodium Acetate buffer (50mM) without any salt, followed by sonication and centrifugation. The soluble fraction containing vUb was loaded onto the SP FF column (GE), and the protein was eluted at a 300-400mM salt gradient. Eluted fractions were concentrated and injected in a Superdex SD75 16/600 Prep grade (GE) column using AKTA pure machine (GE). The pure fractions of vUb were concentrated and stored at −20°C in 25mM Sodium Acetate (pH 5.0) and 100mM NaCl.

NMR samples of vUb were prepared by growing BL21-DE3* cells in M9 media containing isotopic Ammonium chloride (^15^NH4Cl) or isotopic Glucose (^13^C-Glucose). Protein was purified in the same manner as mentioned above. The final buffer for NMR structural experiments contained 25mM Sodium Acetate (pH 5.0) and 100mM NaCl.

Unlike vUb, K54-vUb mutant protein was insoluble after sonication. Hence the pellet was incubated in 8M urea buffer at 4°C for 12-14 hrs. After centrifugation the unfolded supernatant fraction was kept for serial dialysis from 6M, 3M, 1.5M, 0.75M, 0.375M and finally at 0M, for 4-6hrs each, for gradual refolding of protein. The refolded protein was loaded in SP FF column and eluted at a 300-400mM salt gradient. Finally, size exclusion chromatography was carried out to get the pure protein which was stored at −20°C in 25mM Sodium Phosphate buffer (pH 7.5) and 100mM NaCl.

### Purification UbE1, E2s, E3s, and Ub proteins

UbE1 clone was a generous gift from Dr. Cynthia Wolberger lab, and their respective purification protocol was followed for purifying UbE1 protein (44).

All the E2s, E2d2, E225k, Ubc13, and its cofactor Mms2 used in this study were tagged with 6XHis at their N-terminus. Ube2d2/UbcH5a plasmid was a gift from Wade Harper (Addgene: 15782). All his-tagged proteins were purified using IMAC FF (GE) columns followed by their elution with 200-300mM Imidazole linear gradient. Eluted fractions were loaded on Superdex SD75 16/600 Prep grade column, which was pre-equilibrated in 50mM Tris buffer (pH7.5) and 250mM NaCl, to get the pure protein and finally stored at −20°C.

Purification of Ubiquitin proceeded in the same manner as described for vUb. GST-RNF38 clone was a generous gift from Prof. Danny Huang. Purification of GST-RNF38 and GST-RNF4 was done using GST beads (Protino) and eluted out with 10mM reduced Glutathione. Pure protein was obtained after SEC in 50mM Tris buffer with 250mM NaCl and store at −20°C.

### Purification of Proteasomal receptors

Clones of proteasome receptors Rpn10/S5a (UIM motifs: 196-306aa) and hRpn13 (Pru domain: 1-150aa) were a kind gift from Dr. Kylie Walters lab. Both the clones were tagged with N-terminus 6XHis-tag, and their respective purification was performed as followed for other his tagged E2 proteins mentioned above.

### NMR assignments and Structure Calculations

^13^C, ^15^N-labeled vUb protein in 25mM Sodium Acetate pH 5.0 buffer with 100mM NaCl was concentrated up to 1mM, dissolved in 90%-10% v/v H_2_O-D_2_O buffer. NMR spectra were recorded at 298K on 600 MHz Bruker Avance III HD spectrometer equipped with a cryo-probe head. 2D ^1^H-^15^N-HSQC and 3D CBCA(CO)NH, HNCACB, HNCA, HN(CO)CA, HNCO experiments were used for backbone assignments. Sidechain assignments were determined by recording 3D H(CCO)NH, (H)CC(CO)NH and HCCH-TOCSY experiments. 2D (HB)CB (CGCD)HD experiment data along with 2D ^13^C aromatic HSQC spectra were used for assigning aromatic resonances. Standard 3D ^15^N-NOESY-HSQC and ^13^C-NOESY-HSQC experiments with a mixing time of 100ms were used to get NOE restraints. All NMR data were processed in NMRpipe (45) and analyzed by NMRFAM-SPARKY software (19). Following peak-picking of the backbone and side-chain experimental data in SPARKY, the peaks were assigned by PINE and confirmed manually. Automatic peak picking was performed for NOESY spectra. NOESY peak assignments were performed in CYANA (20) and corrected over multiple cycles of structure building. TALOS+ (18) was used to predict the phi and psi torsion angles from the assigned chemical shifts of backbone atoms. Phi/psi torsion angles and NOESY based distance restraints were used to determine the solution structure of vUb by CYANA software. 100 starting structures were calculated, and out of which 20 lowest energy structure models were chosen to represent the final structural ensemble. The backbone dihedral angles of the final converged structures were evaluated by the Molprobity (21) and PSVS (22) suite of programs. Details of the NMR restraints and structure evaluation are provided in Table 1.

### NMR Dynamics

For NMR dynamics studies, uniformly labeled ^15^N-vUb protein (500µM) was dialyzed in 20mM Sodium phosphate buffer, 100mM NaCl with pH 6.0. All the relaxation datasets were obtained at 298K on Bruker Avance 800MHz instrument equipped with a cryoprobe. Longitudinal (T1), Transverse (T2) time constraints and hetero-Nuclear Overhauser Enhancement (hetNOE) experiments were carried out using standard pulse sequences in Bruker. For T1 measurements, HSQC data were recorded at following relaxation delays: 0.004, 0.03, 0.06, 0.1, 0.15, 0.2, 0.4 and 0.8 sec. For T2 measurements, the relaxation delays were set to 0.002, 0.004, 0.008, 0.016, 0.032, 0.064, 0.096 and 0.128 sec. ^15^N-Heteronuclear NOE values were taken as the ratio of peak intensities observed for experiments with 5 sec of ^1^H-presaturation during the recycle delay and the reference experiment without the presaturation for which the recycle delay was set to 10sec. Order parameter (S^2^) was calculated using T1, T2, and hetNOE experiments and then fitted to spectral density function (23, 24) in Bruker Dynamic Centre software. The error of the fits was generated using Monte Carlo simulation.

### Denaturation Melts

#### a. Chemical Melt

40µM of protein was incubated overnight in 20mM Sodium Phosphate buffer (pH 6) containing 100mM Salt and different concentrations of guanidium hydrochloride from 0-6M at room temperature. The unfolding of protein was probed by monitoring far-UV CD signal at 222nm using Jasco J-815 spectropolarimeter.

The data was analyzed and fitted to a two-state (N→U) unfolding process using Sigma plot software where the raw data was converted into fraction unfolded (Fu) values using the following equation (46):

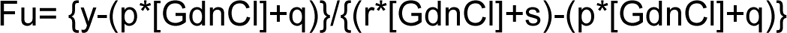

Here, y is the CD value at a particular concentration of [GdnCl]; p and r represent slopes whereas q and s represent intercepts of the native and unfolded protein baselines respectively. Further fitting the curve of Fu vs. [GdnCl] using monomeric 2-para melt equation, the free energy of unfolding, ΔG_unfolding_, and slope of the transition curve, m, were obtained. Finally, the free energy (ΔG_0_) at 0M [GdnCl] is calculated by fitting ΔG_unfolding_ and m in the following 2-state unfolding equation (47):

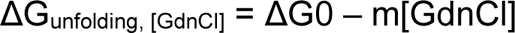

#### b. Thermal Melt

40µM of protein in Sodium Phosphate buffer and 100mM Salt was used for thermal melt. The mean residual ellipticity at 222nm was measured using Jasco J-815 spectropolarimeter, which is connected to a Peltier. The data were recorded from 30°C to 95°C after every 1°C per minute rise in temperature with 32s of data integration time and finally fitted with a two-state folding curve to obtain Tm.

### Temperature Coefficient

Temperature coefficients experiments were performed using uniformly labeled ^15^N-vUb and ^15^N-Ub samples dialyzed in 25mM NaOAc, 50mM NaCl, and pH 5.8. A total of 9 HSQC spectra were collected for both the proteins from 283K to 323K after every 5K increment. All the NMR HSQC experiments were processed by NMRPipe and analyzed in Sparky. Only shifts in ^1^H-dimension were considered and later analyzed using linear regression by MATLAB. The slope of each plot between ΔδNH and temperature is termed as temperature coefficients (ΔδNH/ΔT).

### Binding with Proteasome receptors

#### NMR titrations

For NMR titration experiments, 110 µM ^15^N-vUb was titrated with 700 µM of S5a at following molar ratios 0.25, 0.5, 1, 2, 3, 4 and 5. The reverse titration was performed using and 105 µM ^15^N-S5a and 1.3 mM vUb in 0.25, 0.5, 1, 2, 3, 4, and 5 molar ratios. Both the titration was performed in 20 mM Sodiµm Phosphate buffer (pH 6) with 100 mM NaCl. Chemical shift assignments of S5a was taken from BMRB server (ID: 6233). All the titration data were processed in NMRpipe and analyzed in Sparky software. Using MATLAB, the titration data was fit in either 1:1 or 1:2 protein: ligand model using the equation CSP_obs_ = CSP_max_ {([P]_t_+[L]_t_+K_d_) - [([P]_t_+[L]_t_+K_d_)^2^-4[P]_t_[L]_t_]^1/2^}/2[P]_t_, where [P]t and [L]t are total concentrations of protein and ligand at any titration point.

NMR titration experiments for 15N-vUb (100uM) and hRpn13 (400uM) were performed at following molar ratios 0.5, 1, 2, 3 and 4 in 20 mM Sodiµm Phosphate buffer (pH 7.5) with 250mM NaCl. However, due to intermediate exchange and high affinity, the K_d_ values could not be calculated between vUb and Rpn13.

#### ITC Binding Assay

vUb and S5a were co-dialyzed in 20mM sodium phosphate buffer (pH 7.5) containing 100mM NaCl. 4mM vUb was injected into the sample cell containing 80µM of S5a at 25°C. The measurement was performed on a Microcal ITC200 (GE Healthcare), and binding isotherm was plotted and analyzed using Origin (v7.0). Integrated interaction heat values from individual experiments were normalized, and the data were fit, omitting the first point, using a single binding site model.

#### Affinity measurements by fluorescence spectroscopy

Change in the intrinsic fluorescence of Tryptophan in Rpn13 upon addition of vUb was monitored, and hence the K_d_ value was determined. The intrinsic fluorescence emission scan from 300nm to 400nm was measured using Horiba Fluromax-4 after excitation at 295nm. The slit width for excitation and emission was kept at 3nm. The decrease in fluorescence was observed after adding vUb with increasing molar quantities from 0.5nM till 10uM. The change in fluorescence intensity at 340nm at different vUb molar ratios was used to generate the binding curve (Fig 5H). The total bound fraction was calculated and normalized and then fitted to one-site specific binding curve in Graph pad Prism to calculate K_d_.

#### *In-vitro* Ubiquitination

*In-vitro* ubiquitination reactions were carried out using 1µM of UbE1, 5µM of E2s, 10µM of E3s and 50µM of Ub and vUb. The reaction was activated by reaction buffer containing 50mM Tris (pH 7.5), 100mM NaCl, 2.5mM MgCl2 and 5mM ATP and incubated for 1hr at 37°C. The ubiquitination reaction was quenched with 25mM EDTA and further ran on 15% SDS gel. For probing poly-ubiquitin chains, immunoblotting was performed using Ubiquitin monoclonal antibody (P4D1 from ENZO) at 1:5000 dilution and HRP conjugated anti-mouse secondary antibody at 1:10000 dilutions. The chemiluminescence was observed by clarity ECL (Bio-Rad) staining in Image Quant LAS 4000 (GE). The time point based assay was carried out with 5µM of Ub and vUb. The SDS-PAGE gel was stained overnight with SYPRO® Ruby protein gel stain and imaged in UVITEC Cambridge.

#### *In-vitro* Deubiquitination Assay

For the Deubiquitination assays, 5µl of Ub and vUb and 10µl of vUb-K54 of the quenched ubiquitination reaction mixture, which was carried out by E2D2 and RNF38, was incubated with different DUBs such as USP2_CD_, OTUBAIN, and GST-AMSH. USP2_CD_ and OTUBAIN were activated in DUb activation buffer (25mM Tris, 150mM NaCl, and 10mM DTT) for 15 minutes at RT before initiating their respective deubiquitination reactions. The deubiquitination reaction was carried out at 37°C for 1hr, quenched with SDS loading dye and immunoblotted onto PVDF membrane and finally probed using Ubiquitin antibody (as mentioned above).

#### Mass Spectrometry

In-vitro ubiquitination reaction of vUb and K54-vUb was carried out using UbE1 (0.5µM), E2D2 (10µM) and RNF-38 (10µM) for 1.5hrs at 37°C. The reaction mixture (100µl) was quenched with SDS-loading dye and loaded in 15% SDS-PAGE. Band approximating 16kDa from vUb and K54-vUb were cut out and given for MS/MS analysis. The gel pieces of di-vUb and di-vUb-K54 were washed with LC-MS grade water for 20minutes followed by vortexing in 400µl of 100mM Ammonium bicarbonate/Acetonitrile [1:1 v/v], till the gel pieces became transparent. 400µl of Acetonitrile was then added and incubated at room temperature with occasional vortexing until the gel pieces became white and shrunken. For trypsin digestion, 100µl of trypsin buffer (13ng/µl of trypsin in 10mM Ammonium bicarbonate containing 10% acetonitrile) was added to dried gel pieces and kept for 2hrs at −20°C. 100µl of Ammonium bicarbonate buffer was added to wet the gel pieces, and the tubes were incubated overnight at 37°C for trypsin cleavage. 5% formic acid/Acetonitrile was added to the solution and incubated for 15mins at 37°C to inactivate trypsin. The gel pieces were carefully removed, and the supernatant was centrifuged for 20minutes before lyophilization and then finally re-solubilized in 40µl of 0.1%formic acids, 2% [v/v] acetonitrile for LCMS analysis. EASY SPRAY PEPMAP RSLC C18 analytical column was used to separate the peptides by the gradient developed from 0.1% [v/v] formic acid to 80% [v/v] Acetonitrile, 0.1% [v/v] formic acid in HPLC grade water over 53mins at a flow rate of 300nL/min in EASY-nLC 1200 system which is in-line coupled to nano-electrospray source of a LTQ-Orbitrap Discovery hybrid mass spectrometer (Thermo Scientific, SanJose, CA, USA). Full MS in the mass range between m/z 375 and m/z 1700 was performed on an Orbitrap Mass analyzer, followed by CID-based MS/MS, performed in the scan range of m/z 100 and m/z 2000. Mascot Distiller software was used to identify the peptides with GG modification, in which the peptide mass tolerance threshold was set to 10ppm, and the maximum fragment mass tolerance was set between 0.2-0.6Da.

### Cell culture experiments

1XFLAG-Ub, 1XFLAG-vUb, and 1XFLAG-vUb-K54 were cloned in the pcDNA3.2 vector. Approximately 50,000 cells were seeded in each well of 12 well plates and incubated for 12hrs at 37°C with 5% CO_2_, before transfecting them with 1XFLAG-Ub, 1XFLAG-vUb and 1XFLAG-vUb-K54 respectively using Lipofectamine 3000 reagent and further, the cells were incubated for 24hrs. The cells were then treated with MG132 for 12hrs before lysing the cells in 1XTBS containing 0.5% NP40 and 1X protease inhibitor cocktail. Total protein was estimated using BCA reagent (Thermo), and the final amount of 3µg of FLAG-Ub and 15µg of FLAG-vUb and FLAG-vUbK54 were loaded on 10%SDS-PAGE gel. Immunoblots were probed with Flag antibody.

## Acknowledgments

This work was supported by intramural grants from the Tata Institute of Fundamental Research. We thank Dipendra Basu for helping in making ML Tree. All the NMR spectra were collected at NMR Facility, and Mass spec data were collected at proteomics unit of Mass Spectrometry facility of National Centre for Biological Sciences. H. N. is a recipient of fellowship from the University Grants Commission. R.D. is a recipient of a DBT-Ramalingaswamy fellowship (BT/HRD/23/02/2006).

## Conflict of Interest

The authors declare that they have no conflict of interest with the content of this article.

## Databank submission

The atomic coordinates and structure factors (PDB id: 6KNA) have been deposited in the Protein data bank (http://wwpdb.org/). The NMR chemical shift is deposited in the Biological Magnetic Resonance Bank under accession number 36278.

## References

1. Yau, R., and Rape, M. (2016) The increasing complexity of the ubiquitin code. Nat. Cell Biol. 18, 579–586

2. Swatek, K. N., and Komander, D. (2016) Ubiquitin modifications. Cell Res. 26, 399–422

3. Varshavsky, A. (2017) The Ubiquitin System, Autophagy, and Regulated Protein Degradation. Annu. Rev. Biochem. 86, 123–128

4. Komander, D., and Rape, M. (2012) The Ubiquitin Code. Annu. Rev. Biochem. 81, 203– 229

5. Hofmann, R. M., and Pickart, C. M. (1999) Enzyme Functions in Assembly of Novel Polyubiquitin Chains for DNA Repair. Cell. 96, 645–653

6. Locke, M., Toth, J. I., and Petroski, M. D. (2014) Lys 11 - and Lys 48 -linked ubiquitin chains interact with p97 during endoplasmic-reticulum-associated degradation. Biochem. J. 459, 205–216

7. Viswanathan, K., Klaus, F., and DeFilippis, V. (2010) Viral hijacking of the host ubiquitin system to evade interferon responses. Curr. Opin. Microbiol. 10.1016/j.mib.2010.05.012

8. Isaacson, M. K., and Ploegh, H. L. (2009) Ubiquitination, Ubiquitin-like Modifiers, and Deubiquitination in Viral Infection. Cell Host Microbe. 5, 559–570

9. Wimmer, P., and Schreiner, S. (2015) Viral Mimicry to Usurp Ubiquitin and SUMO Host Pathways. Viruses. 10.3390/v7092849

10. Boutell, C., Everett, R., Hilliard, J., Schaffer, P., Orr, A., and Davido, D. (2008) Herpes Simplex Virus Type 1 ICP0 Phosphorylation Mutants Impair the E3 Ubiquitin Ligase Activity of ICP0 in a Cell Type-Dependent Manner ᰔ. J. Virol. 82, 10647–10656

11. Chaurushiya, M. S., Lilley, C. E., Aslanian, A., Meisenhelder, J., Scott, D. C., Ticau, S., Boutell, C., Yates, J. R., Schulman, B. A., Hunter, T., and Weitzman, M. D. (2012) Viral E3 Ubiquitin Ligase-Mediated Degradation of a Cellular E3 : Viral Mimicry of a Cellular Phosphorylation Mark Targets the RNF8 FHA Domain. Mol. Cell. 46, 79–90

12. Slack, J., and Arif, B. M. (2007) The Baculoviruses Occlusion-Derived Virus: Virion Structure and Function. in Advances in Virus Research, pp. 99–165, 69, 99–165

13. Ayres, M. D., Howard, S. C., Kuzio, J., Lopez-Ferber, M., and Possee, R. D. (1994) The Complete DNA Sequence of Autographa californica Nuclear Polyhedrosis Virus. Virology. 202, 586–605

14. Guarino, L. A. (1990) Identification of a viral gene encoding a ubiquitin-like protein. Proc. Natl. Acad. Sci. U. S. A. 87, 409–413

15. Guarino, L. A., Smith, G., and Dong, W. (1995) Ubiquitin is attached to membranes of baculovirus particles by a novel type of phospholipid anchor. Cell. 80, 301–309

16. Reilly, L. M., and Guarino, L. A. (1996) The Viral Ubiquitin Gene of Autographa californica Nuclear Polyhedrosis Virus Is Not Essential for Viral Replication. Virology. 218, 243–247

17. Biswas, S., Willis, L. G., Fang, M., Nie, Y., and Theilmann, D. A. (2018) Autographa californica Nucleopolyhedrovirus AC141 (Exon0), a Potential E3 Ubiquitin Ligase, Interacts with Viral Ubiquitin and AC66 To Facilitate Nucleocapsid Egress. J. Virol. 92, 1–21

18. Shen, Y., Delaglio, F., Cornilescu, G., and Ad Bax (2009) TALOS + : a hybrid method for predicting protein backbone torsion angles from NMR chemical shifts. J. Biomol. NMR. 44, 213–223

19. Lee, W., Tonelli, M., and Markley, J. L. (2015) Structural bioinformatics NMRFAM-SPARKY : enhanced software for biomolecular NMR spectroscopy. Bioinformatics. 31, 1325–1327

20. Guntert, P., and Buchner, L. (2015) Combined automated NOE assignment and structure calculation with CYANA. J. Biomol. NMR. 62, 453–471

21. Chen, V. B., Bryan, W., Iii, A., Headd, J. J., Keedy, D. A., Robert, M., Kapral, G. J., Murray, L. W., Jane, S., and David, C. (2010) MolProbity : all-atom structure validation for macromolecular crystallography research papers. Acta Crystallogr. Sect. D. D66, 12–21

22. Vuister, G. W., and Fogh, R. H. (2014) An overview of tools for the validation of protein NMR structures. J. Biomol. NMR. 58, 259–285

23. Lipari, G., and Szabo, A. (1982) Model-Free Approach to the Interpretation of Nuclear Magnetic Resonance Relaxation in Macromolecules. 1. Theory and Range of Validity. J. Am. Chem. Soc. 140, 4546–4559

24. Lipari, G., and Szabo, A. (1982) Model-Free Approach to the Interpretation of Nuclear Magnetic Resonance Relaxation in Macromolecules. 2. Theory and Range of Validity. J. Am. Chem. Soc.

25. Surana, P., and Das, R. (2016) Observing a late folding intermediate of Ubiquitin at atomic resolution by NMR. Protein Sci. 25, 1438–1450

26. Lee, S. Y., Pullen, L., Virgil, D. J., Castañeda, C. A., Abeykoon, D., Bolon, D. N. A., and Fushman, D. (2014) Alanine Scan of Core Positions in Ubiquitin Reveals Links between Dynamics, Stability, and Function. J. Mol. Biol. 426, 1377–1389

27. Sankar Chatterjee, K., Tipathi, V., and Das, R. (2019) A conserved and buried edge-to-face aromatic interaction in small ubiquitin-like modifier (SUMO) has a role in SUMO. J. Biol. Chem. 294, 6772–6784

28. Peña, A. H. De, Goodall, E. A., Gates, S. N., Lander, G. C., and Martin, A. (2018) Substrate-engaged 26 S proteasome structures reveal mechanisms for ATP-hydrolysis – driven translocation. Science (80-.). 362, 1018

29. Deverauxf, Q., Ustrellf, V., Pickart, C., and Rechsteiner, M. (1994) A 26 S Protease Subunit That Binds Ubiquitin Conjugates*. J. Biol. Chem. 269, 7059–7061

30. Husnjak, K., Elsasser, S., Zhang, N., Chen, X., Randles, L., Shi, Y., Hofmann, K., Walters, K. J., Finley, D., and Dikic, I. (2008) Proteasome subunit Rpn13 is a novel ubiquitin receptor. Nature. 453, 481–488

31. Wang, Q., Young, P., Walters, K. J., and Hall, J. (2005) Structure of S5a Bound to Monoubiquitin Provides a Model for Polyubiquitin Recognition. J. Mol. Biol. 348, 727–739

32. Low, P., Hastings, R., Dawson, S., Sass, M., Billett, M., Mayer, R., and Reynolds, S. (2000) Localisation of 26S proteasomes with different subunit composition in insect muscles undergoing programmed cell death. Cell Death Differ. 7, 1210–1217

33. Zhang, N., Wang, Q., Ehlinger, A., Randles, L., Lary, J. W., Kang, Y., Haririnia, A., Storaska, A. J., Cole, J. L., Fushman, D., and Walters, K. J. (2009) Structure of the S5a : K48-Linked Diubiquitin Complex and Its Interactions with Rpn13. Mol. Cell. 35, 280–290

34. Hofmann, R. M., and Pickart, C. M. (2001) In Vitro Assembly and Recognition of Lys-63 Polyubiquitin Chains *. J. Biol. Chem. 276, 27936–27943

35. Middleton, A. J., and Day, C. L. (2015) The molecular basis of lysine 48 ubiquitin chain synthesis by Ube2K. Sci. Rep. 5, 1–14

36. Singh, R. K., Kazansky, Y., Wathieu, D., and Fushman, D. (2017) Hydrophobic Patch of Ubiquitin is Important for its Optimal Activation by Ubiquitin Activating Enzyme E1. Anal. Chem. 89, 7852–7860

37. Burch, T. J., and Haas, A. L. (1994) Site-Directed Mutagenesis of Ubiquitin. Differential Roles for Arginine in the Interaction with Ubiquitin-Activating Enzyme ? Biochemistry. 33, 7300–7308

38. Baboshina, O. V, and Haas, A. L. (1996) Novel Multiubiquitin Chain Linkages Catalyzed by the Conjugating Enzymes E2 EPF and RAD6 Are Recognized by 26 S Proteasome Subunit 5 *. J. Biol. Chem. 271, 2823–2831

39. Ma, S. S., Zhang, Z., Xia, H. C., Chen, L., Yang, Y. H., Yao, Q., and Chen, K. P. (2015) Evolutionary analysis of the ubiquitin gene of baculovirus and insect hosts. Genet. Mol. Res. 14, 9963–9973

40. Kumar, S., Stecher, G., Li, M., Knyaz, C., and Tamura, K. (2018) MEGA X : Molecular Evolutionary Genetics Analysis across Computing Platforms. 35, 1547–1549

41. Lanfear, R., Frandsen, P. B., Wright, A. M., Senfeld, T., and Calcott, B. (2018) PartitionFinder 2 : New Methods for Selecting Partitioned Models of Evolution for Molecular and Morphological Phylogenetic Analyses. Mol. Biol. Evol. 34, 772–773

42. Stamatakis, A. (2014) RAxML version 8 : a tool for phylogenetic analysis and post-analysis of large phylogenies. Bioinformatics. 30, 1312–1313

43. Miller, M. A., Pfeiffer, W., and Schwartz, T. (2010) Creating the CIPRES Science Gateway for Inference of Large Phylogenetic Trees. Proc. Gatew. Comput. Environ. Work.

44. Berndsen, C. E., and Wolberger, C. (2011) A spectrophotometric assay for conjugation of ubiquitin and ubiquitin-like proteins q. Anal. Biochem. 418, 102–110

45. Delaglio, F., Grzesiek, S., Vuister, G. W., Zhu, G., Pfeifer, J., and Ad Bax (1995) NMRPipe : A multidimensional spectral processing system based on UNIX pipes *. J. Biomol. NMR. 6, 277–293

46. Privalov, P. (1979) Stability of Proteins. in Advances in Protein Chemistry, pp. 167–241

47. Agashe, V. R., and Udgaonkar, J. B. (1995) Thermodynamics of Denaturation of Barstar: Evidence for Cold Denaturation and Evaluation of the Interaction with Guanidine Hydrochloride+. Biochemistry. 34, 3286–3299

