## Supplemental Information for "A novel polyubiquitin chain linkage formed by viral Ubiquitin prevents cleavage by deubiquitinating enzymes"

### Supporting Information

*Electronic address:

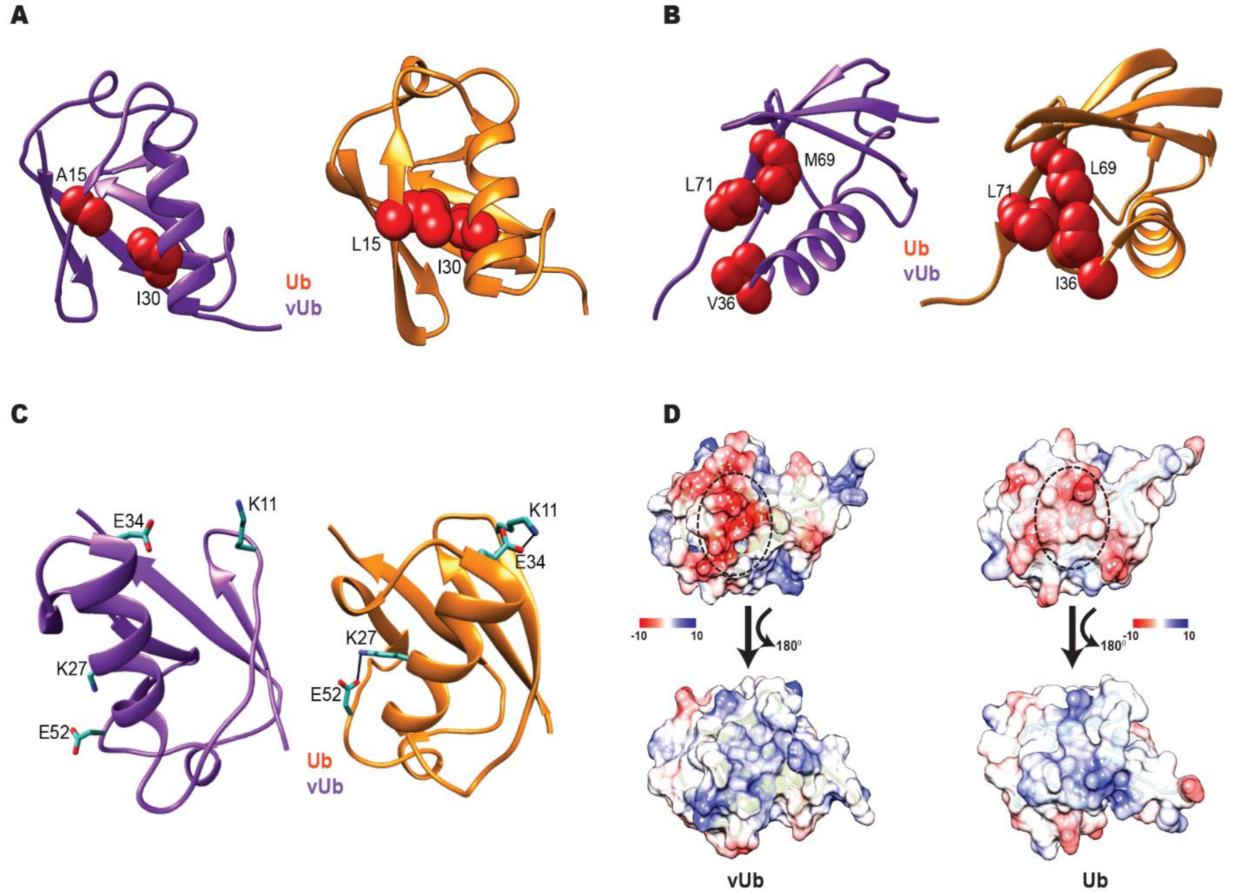


**Supplementary Figure S1**: Structural feature of vUB. (A) The core hydrophobic contacts are compared between Ub and vUb is shown respectively. L15 in β2 forms tight contact with I30 in α1 of Ub. It is altered to a smaller sidechain residue A15 in vUb, and cannot form contact with I30. (B) I36 contacts L69 and L71 of β5 via long hydrophobic side chain in Ub, whereas, this contact is hindered since I36 is substituted to V36 in vUb which is a small side chain containing residue. (C) The two salt bridges K11/E34 and K27/E52 present in Ub are absent in vUb. (D) Color gradient scheme of the electrostatic surface potential of vUb and Ub is shown. Positively, Negatively, and neutral surfaces are shown as blue, red, and white colors, respectively.

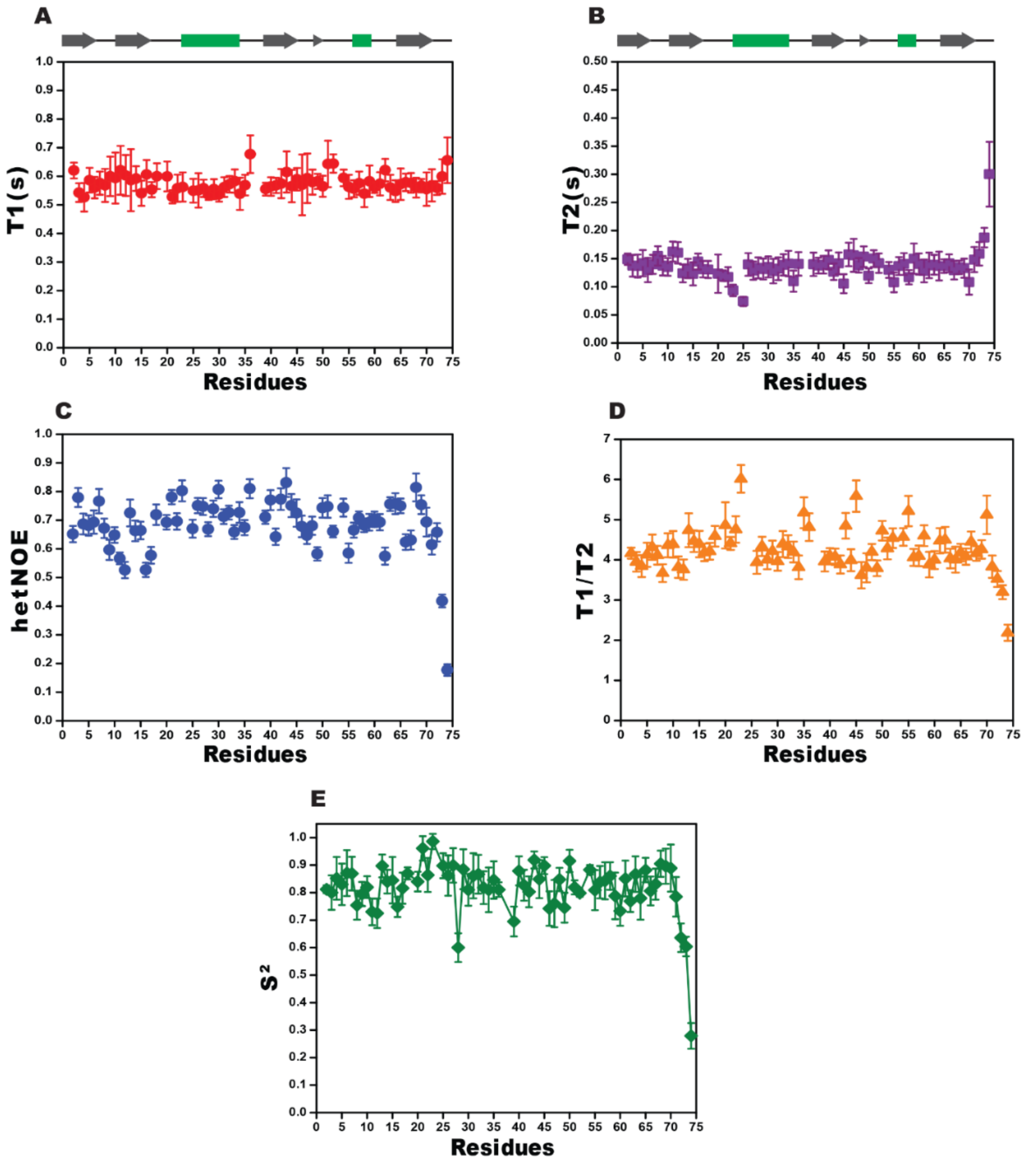


**Supplementary Figure S2**: Relaxation experiments of Ub. (A) T1, (B) T2 and (C) hetNOEs were plotted against residue number of Ub. (D) The T1/T2 ratio was calculated and plotted against the residue number of Ub. (E) Using A, B, and C order parameter ($S^2$) was calculated using the Lipari Szabo model. The secondary structure elements of Ub is provided on top of each plot.

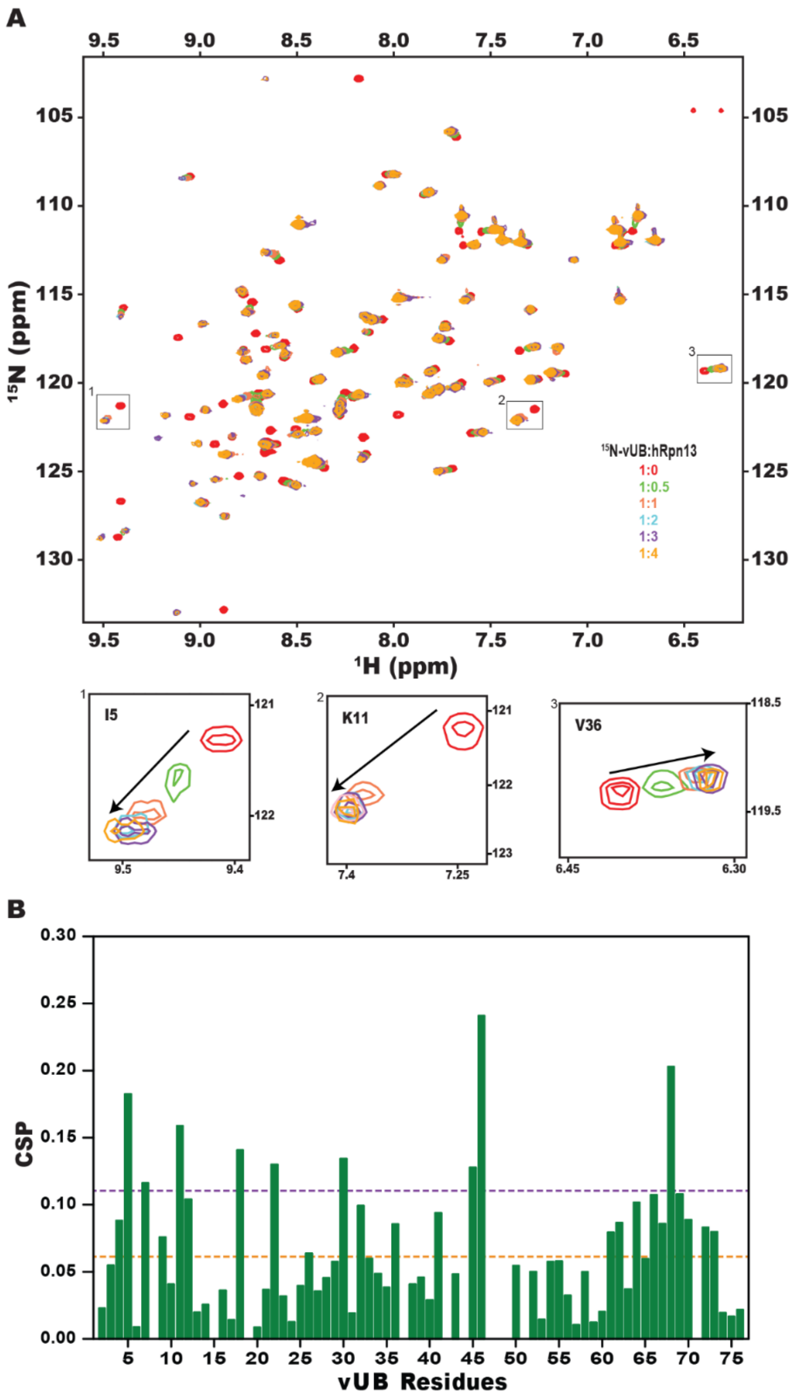


**Supplementary Figure S3**: Binding of vUB with hRpn13. (A) Overlay of $^{15}$N-edited HSQC spectra of $^{15}$N-vUb with different stoichiometric ratios of hRpn13. The resonances of residues I5, K11, and V36 are zoomed for clarity. (B) The CSP observed in vUb upon titration with hRpn13 is plotted against vUb residue number. Mean and mean with standard deviation are highlighted in orange and purple, respectively.

S4A

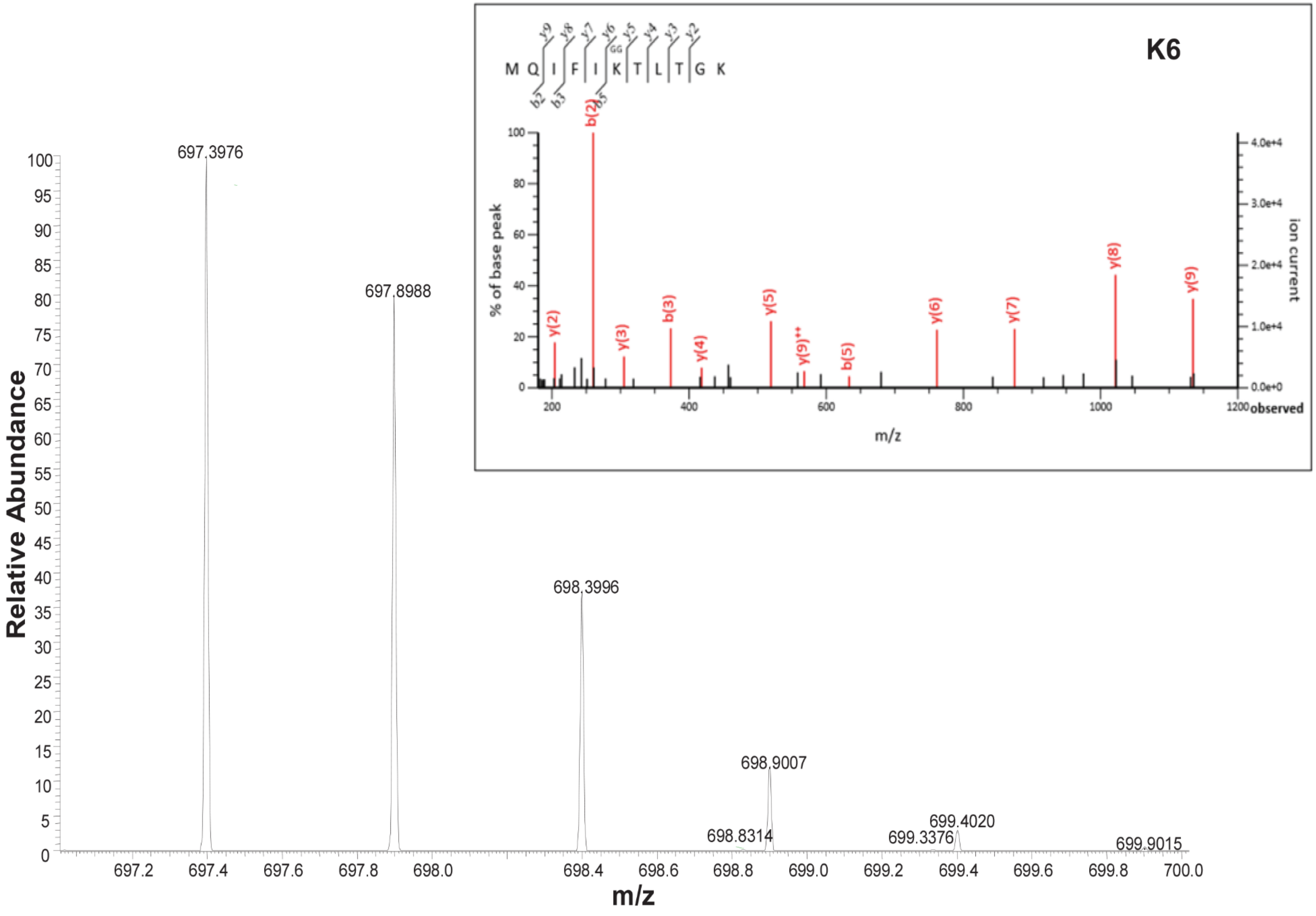

K6
M Q I F I K T L T G K
GG
y9 y8 y7 y6 y5 y4 y3 y2
b2 b3 b5
% of base peak
ion current
observed
m/z
y(2)
b(2)
y(3)
b(3)
y(4)
y(5)
y(9)++
b(5)
y(6)
y(7)
y(8)
y(9)
Relative Abundance
m/z
697.3976
697.8988
698.3996
698.9007
698.8314
699.3376
699.4020
699.9015

S4B

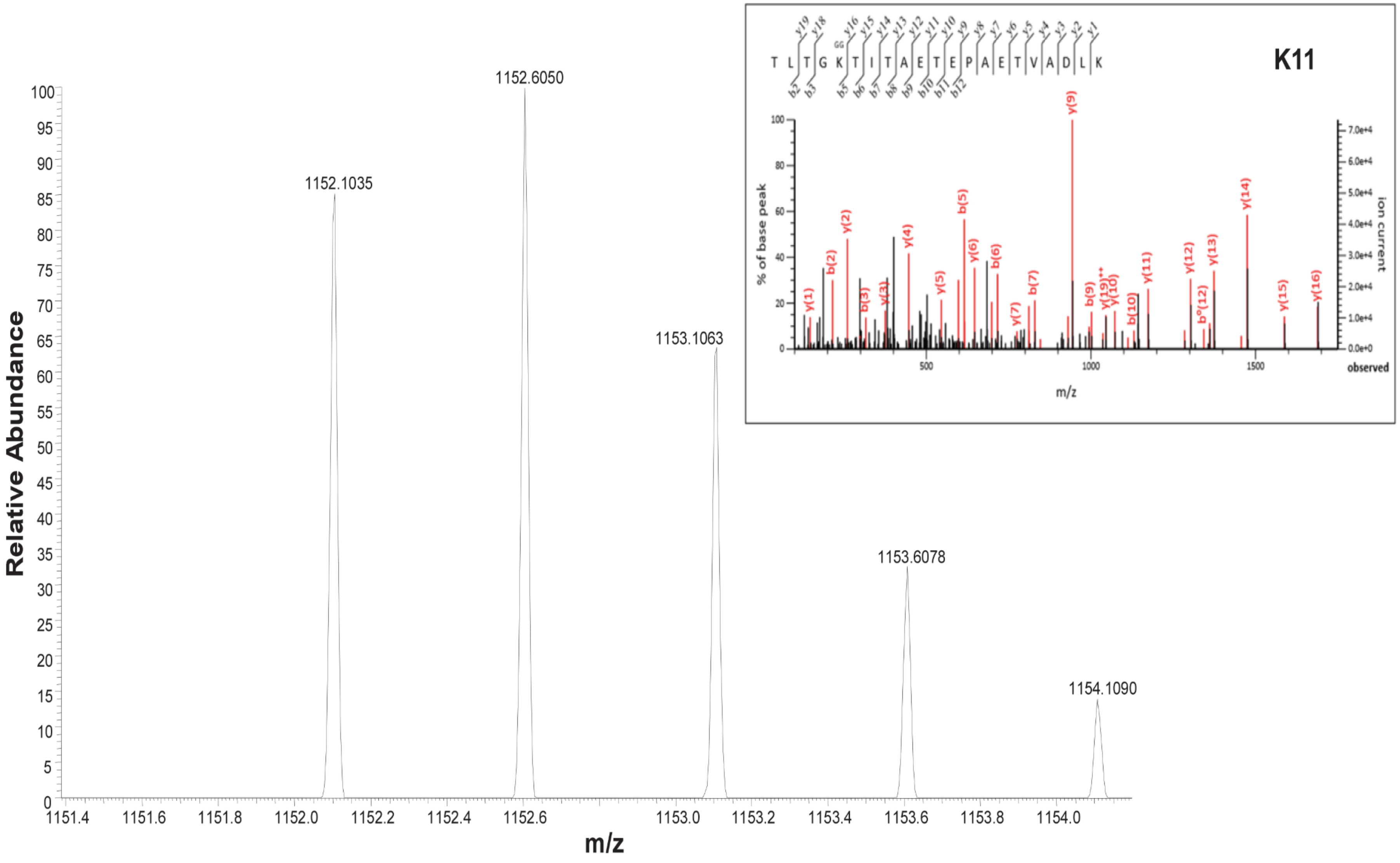

Relative Abundance
m/z
1152.1035
1152.6050
1153.1063
1153.6078
1154.1090
K11
T L T G K T I T A E T E P A E T V A D L K
% of base peak
ion current
observed
y(1) b(2) y(2) b(3) y(3) y(4) y(5) b(5) y(6) b(6) y(7) b(7) y(9) b(9) y(19)++ y(10) b(10) y(11) y(12) b°(12) y(13) y(14) y(15) y(16)

**S4C**

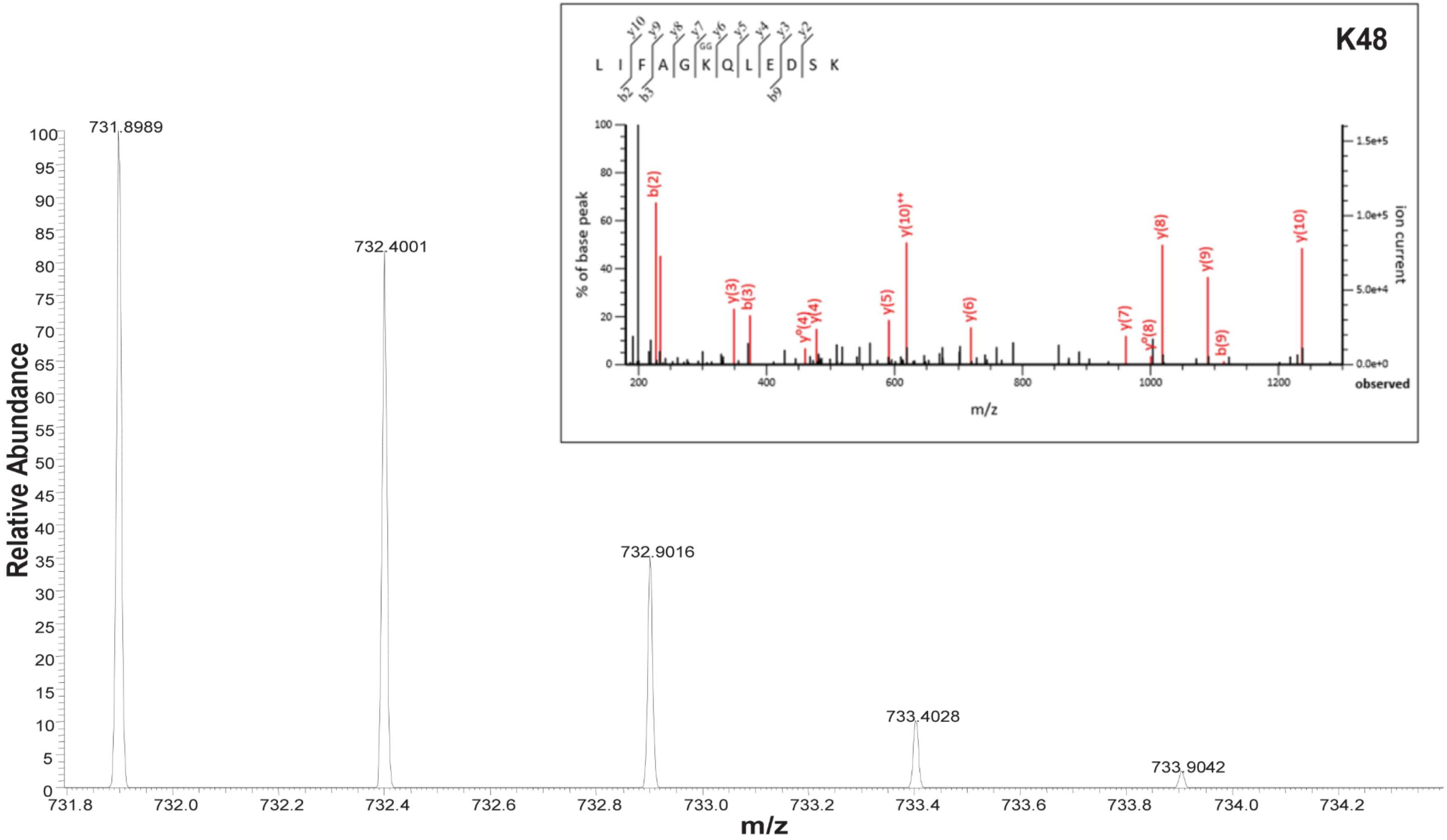
K48
L I F A G K Q L E D S K
GG
y10 y9 y8 y7 y6 y5 y4 y3 y2
b2 b3 b9
% of base peak
ion current
observed
m/z
b(2)
y(3)
b(3)
y°(4)
y(4)
y(5)
y(10)++
y(6)
y(7)
y°(8)
y(8)
y(9)
b(9)
y(10)
1.5e+5
1.0e+5
5.0e+4
0.0e+0
Relative Abundance
731.8989
732.4001
732.9016
733.4028
733.9042
m/z

S4D

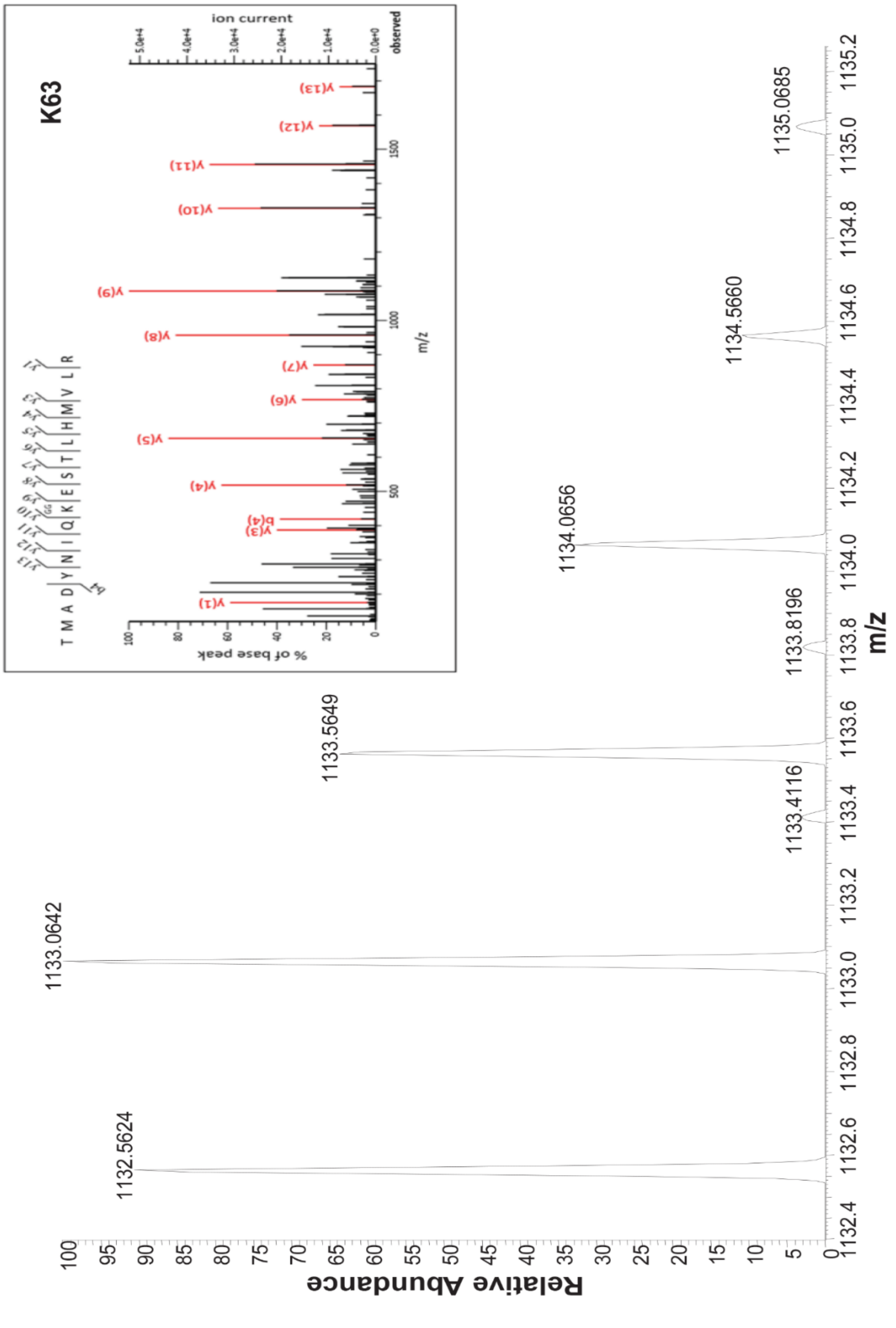


**Supplementary Figure S4**: **A-D** MS-MS spectrum of all the remaining 4 out of 5 tryptic peptides mentioned in Table 4 which shows multiple types of isopeptide linkages are formed in vUB when E2D2/RNF38 are used to carry out polyubiquitination assay.

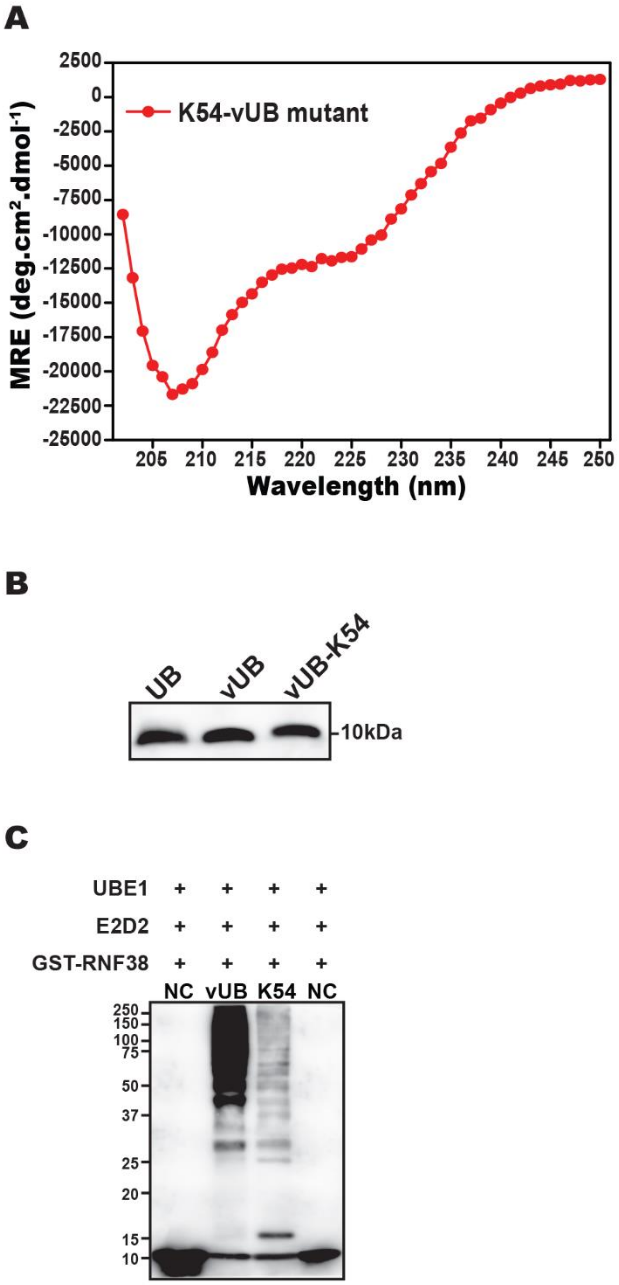


**Supplementary Figure S5**: (A) Far-UV Circular dichroism profile of refolded K54-vUB mutant. The raw data were converted to Mean Residual Ellipticity (MRE) and plotted against wavelength from 250 nm to 202nm. (B) Ub, vUb and K54-vUb was blotted with the same anti-Ub. (C) Polyubiquitination of vUb and K54-vUb by the E2D2/RNF38 complex.

**Table S1: Details of the viral Ubiquitin sequences from the respective 39 insect viral genomes and one Ubiquitin gene sequence from the insect host genome**

| S/N | Virus name | Gene ID | NCBI Reference Sequence |
|---|---|---|---|
| 1 | Adoxophyes honmai NPV | 1485767 | NC_004690.1 |
| 2 | Adoxophyes orana NPV | 11040347 | NC_011423.1 |
| 3 | Lymantria xylina NPV | 8919476 | NC_013953.1 |
| 4 | Lymantria dispar MNPV | 1488589 | NC_001973.1 |
| 5 | Phthorimaea operculella GV | 949328 | NC_004062 |
| 6 | Plutella xylostella GV | 912144 | NC_002593.1 |
| 7 | Xestia c-nigrum GV | 1442286 | NC_002331.1 |
| 8 | Helicoverpa armigera GV | 10973731 | NC_010240 |
| 9 | Pseudaletia unipuncta GV | 8763910 | NC_013772.1 |
| 10 | Choristoneura fumiferana GV | 4155894 | NC_008168.1 |
| 11 | Artogeia rapae GV | 11107052 | NC_013797.1 |
| 12 | Cydia pomonella GV | 921351 | NC_002816.1 |
| 13 | Trichoplusia ni single NPV | 5141966 | NC_007383.1 |
| 14 | Chrysodeixis chalcites NPV | 3431485 | NC_007151.1 |
| 15 | Clanis bilineata NPV | 5141827 | NC_008293.1 |
| 16 | Spodoptera litura NPV | 922137 | NC_003102.1 |
| 17 | Hyphantria cunea NPV | 3890584 | NC_007767.1 |
| 18 | Antheraea pernyi MNPV-L2 | | EF207986.1$^{+}$ |
| 19 | Antheraea pernyi NPV | 5141439 | NC_008035.3 |
| 20 | Choristoneura fumiferana MNPV | 1482695 | NC_004778.3 |
| 21 | Orgyia pseudotsugata MNPV | 912026 | NC_001875.2 |
| 22 | Epiphyas postvittana NPV | 921803 | NC_003083.1 |
| 23 | Bombyx mori NPV | 1724483 | NC_001962.1 |
| 24 | Maruca vitrata MNPV | 4643040 | NC_008725.1 |
| 25 | Autographa californica NPV | 1403867 | NC_001623.1 |
| 26 | Anticarsia gemmatalis NPV | 5141078 | NC_008520.2 |
| 27 | Choristoneura fumi. DEF MNPV | 2943789 | NC_005137.2 |
| 28 | Helicoverpa armigera NPV-G4 | 920057 | NC_002654.2 |
| 29 | Helicoverpa armigera NPV | 922002 | NC_003094.2 |
| 30 | Ecotropis obliqua NPV | 5176494 | NC_008586.1 |
| 31 | Orgyia leucostigma NPV | 5850470 | NC_010276.1 |
| 32 | Euproctis pseudoconspersa NPV | 7804603 | NC_012639.1 |
| 33 | Spodoptera litura NPV-II | 7057161 | NC_011616.1 |
| 34 | Spodoptera exigua NPV | 2715738 | NC_002169.1 |
| 35 | Spodoptera frugiperda MNPV | 5176125 | NC_009011.2 |
| 36 | Mamestra configurata NPV-A | 935840 | NC_003529.1 |
| 37 | Mamestra configurata NPV-B | 951821 | NC_004117.1 |
| 38 | Agrotis segetum NPV | 3974411 | NC_007921.1 |
| 39 | Agrotis ipsilon MNPV | 6965910 | NC_011345.1 |
| 40 | Bombyx Mandarina* | | DQ839401.1$^{+}$ |

+ Only GenBank iD; ∗ Insect Host (outgroup)
